# Type I interferon signaling instills divergent metastatic phenotypes and immunotherapy responses

**DOI:** 10.1101/2025.09.18.677123

**Authors:** Cort B. Breuer, Marcos Labrado, Grayson E. Rodriguez, K. Christopher Garcia, Nathan E. Reticker-Flynn

**Author notes:** These authors contributed equally (Marcos Labrado, Grayson E. Rodriguez). Lead Contact: **Error! Hyperlink reference not valid.**.

## Abstract

Metastatic colonization of distant tissues is responsible for most cancer deaths, yet while this colonization is typically preceded by lymph node (LN) metastasis, the clonal and functional relationships between metastases in these tissues remain unclear. Reconstruction of metastatic phylogenies in mouse models and patients suggests that LN and distant organ metastases are often seeded by independent clones, yet no genetic alterations have bene found to account for this divergent seeding. Here, we discover opposing roles for interferon signaling in metastasis to LNs and distant organs. We uncover that exposure to leukocyte-derived type I interferon promotes LN metastasis while impairing spread to distant organs by inducing interferon-stimulated gene (ISG) expression. Further, while LN metastases have previously been demonstrated to disrupt anti-tumor immunity, we find that ISG expression induced during LN metastasis sensitizes tumors to immune checkpoint blockade (ICB). By mimicking the phenotype of LN metastases through delivery of type I interferon, ICB refractory primary tumors and distant metastases can be sensitized to ICB. Collectively, these findings demonstrate that the immune system conditions tumors for LN metastasis through type I interferon secretion while impairing distant spread and that this axis can be exploited to render immunotherapy resistant tumors sensitive to immune checkpoint blockade therapy.

Graphical Abstract

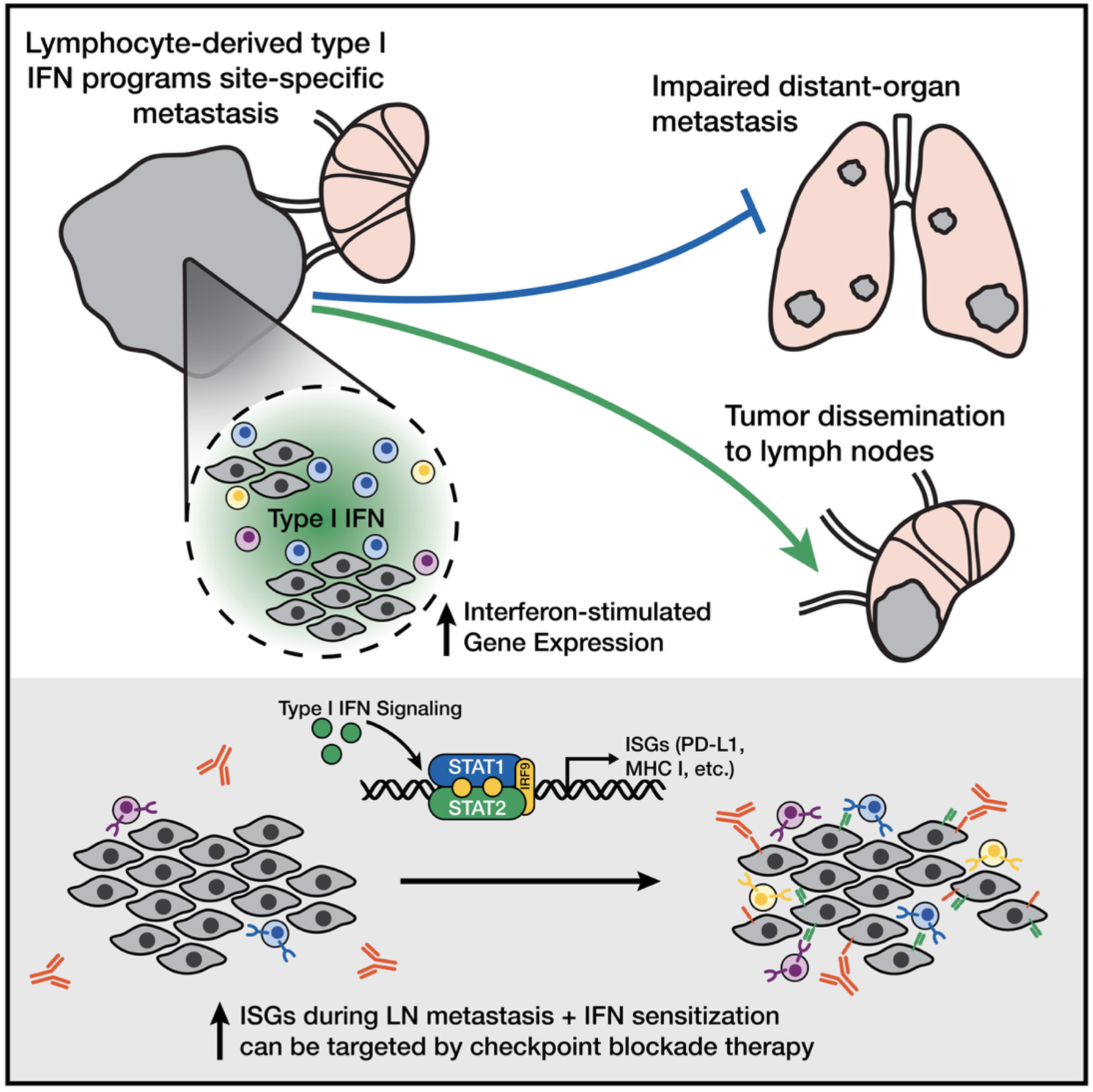

## Introduction

Most cancer deaths are due to metastatic spread rather than primary tumor growth^1,2^. Lymph node (LN) metastasis frequently precedes spread to more distant sites, conferring worse prognosis across most solid cancer types (Supp. Fig. 1A-C)^3–5^, yet the relationships between LN metastases and those in distant tissues remains unclear. The antecedent observation of nodal involvement originally led physicians and scientists to posit that LNs serve as staging grounds for further metastatic dissemination^6–8^. Nonetheless, while tumors within LNs can seed distant sites of metastasis, studies employing phylogenetic reconstruction of metastatic clones by sequencing in mice and humans have suggested that LN metastases are often derived from independent clones within the primary tumor^9–17^. Yet, even though LN and distant organ metastases are often seeded by independent clones, there are no conserved driver gene mutations associated with seeding pattern^18,19^. Furthermore, the elevated polyclonality of LN metastatic seeding with respect to other metastases has led some researchers to suggest that fewer selective pressures are involved in LN metastasis, and consequently, fewer adaptations are required for the acquisition of LN metastatic potential^19–21^. Thus, whether the divergent clonal architecture of LN and distant metastases reflects distinct evolutionary pressures or an absence of selection within LNs remains to be explored.

LNs are essential immune regulatory sites; by acting as collection depots for lymphatic drainage and facilitating immune cell communication, they broadly coordinate both local and systemic immune responses^22–24^. Their central roles in orchestrating immunity, including against tumors, renders their early and frequent metastatic colonization perplexing. We and others recently discovered that by colonizing LNs, tumors induce immune tolerance in a manner that facilitates distant metastasis and relies upon the formation of tumor-specific regulatory T cells (Tregs)^25,26^. It remains unclear what signal initiates LN metastasis and whether this cue solely enhances spread to local LNs or directs cells away from distant sites as would be hypothesized from the divergent seeding patterns previously observed. Further, LN metastases are not solely associated with impaired immunity to metastases but also the generation of dysfunctional immune responses to primary tumors and subsequent immune checkpoint blockade (ICB)^26,27^. The exact role of the tumor-draining LN (tdLN) for ICB response prior to and after LN metastasis remains unclear. Studies in mice perturbing LNs in combination with high-dimensional immune profiling^28–30^, human de-escalation trials^31^, and successful neoadjuvant ICB trials in patients with locally-advanced resectable disease^32–34^, would suggest that tdLNs are necessary sites of ICB response and should be preserved^27,35^. In contrast, assessment of nodal involvement for staging purposes and suggestions that LN metastasis induces dysfunctional states that cannot be rescued by therapy, continue to motivate extensive lymphadenectomy for multiple malignancies^27,36^. Functional assessment of the impact of LN metastasis on ICB response has not been completed to date, leaving open the question of whether T cells in tumor-involved LNs can still mount productive responses to therapy.

Here, we investigate the cues that direct LN metastasis and their impact on distant spread as well as immunotherapy outcomes. We find that type I interferon programs tumors to metastasize to LNs while impairing their ability to spread to distant sites, contributing to independent colonization of metastases. We further find that although LN metastases enhance dysfunctional T cell states, ISG expression acquired during LN metastasis makes tumors vulnerable to checkpoint blockade therapy and a similar ICB-responsive state can be induced at distant metastatic sites through delivery of type I interferon.

## Results

### Leukocyte-derived type I interferon is associated with lymph node metastasis

To study lymph node metastasis, we employed a model we previously developed by *in vivo* selection of increasingly LN metastatic variants of the B16F0 melanoma cell line, syngeneic in immunocompetent C57BL/6 mice^37^. Following a similar approach as previous groups, we isolated rare LN metastases from mice implanted with B16F0 tumors, derived cell lines, followed by reimplantation and selection for additional LN metastases, repeating the process over nine iterative generations (Fig. 1A)^25,38^. This selection scheme resulted in nearly 300 cell lines, of which we performed RNA-sequencing (RNA-seq) on 32 representative cell lines, spanning all nine generations.

**Figure 1.**
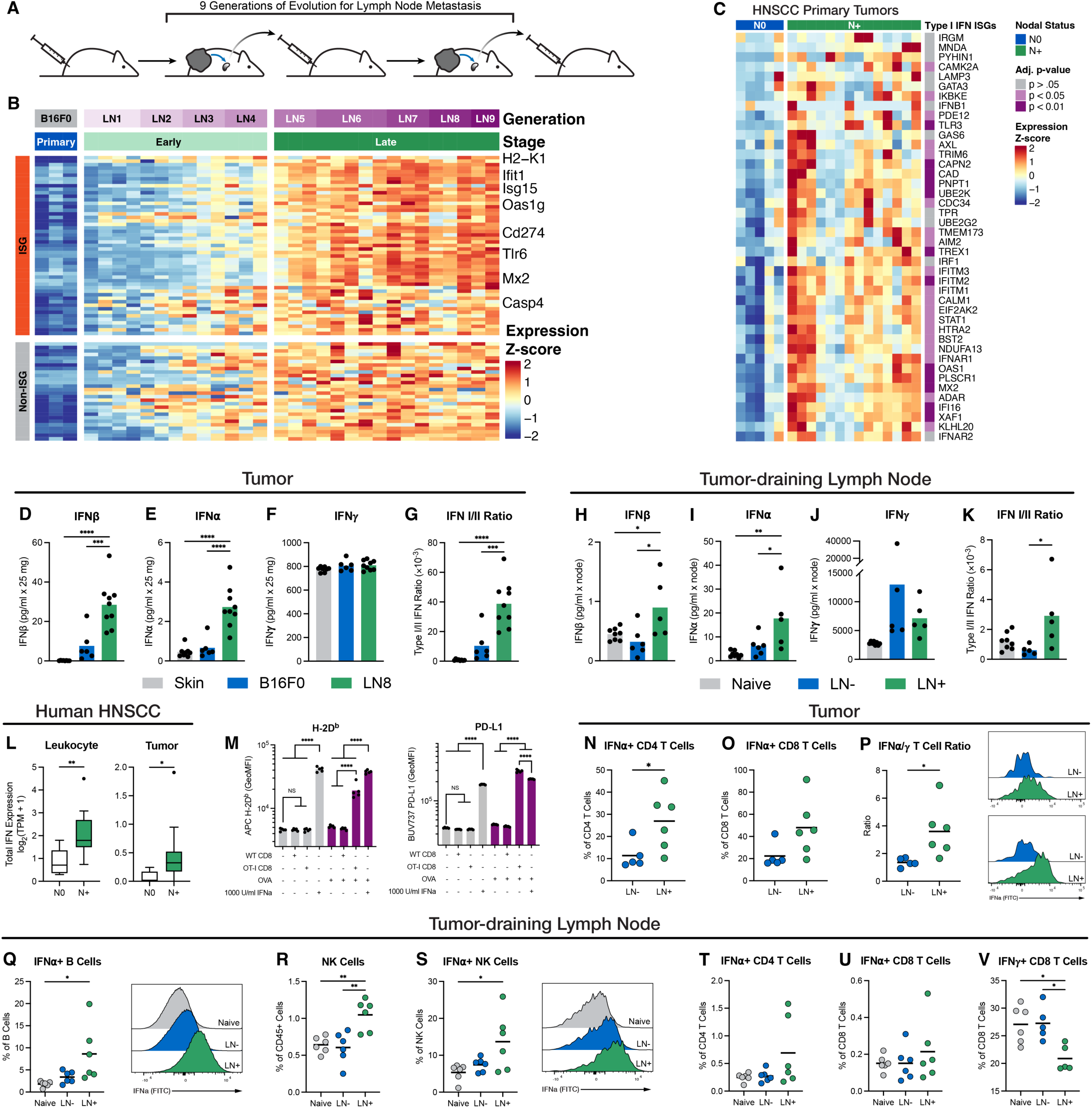
Lymphocyte-derived type I interferon induces ISG expression during LN metastasis. (A) Schematic of evolution of melanoma LN-metastatic cell line model. (B) Heatmap of differentially expressed genes between B16F0 and late stage LN-metastatic cell lines, grouped according to ISG annotation. (C) Heatmap of type I interferon responsive gene-set expression for bulk-sorted tumor cells from HNSCC patient tumors according to nodal involvement. (D-K) Secreted levels of IFNa, IFNb, IFNy, and type I/II interferon ratio for tumor-site or tumor-draining LN for naïve, primary tumor, or LN metastatic mice. (L) Total interferon gene expression scores for bulk-sorted leukocytes or tumor cells from HNSCC primary tumors according to nodal involvement. (M) ISG expression on B16F0 tumor cells during non-specific or antigen-specific T cell co-culture. (N-V) Immune cell frequency and interferon production flow cytometry staining for tumors and tumor-draining LNs from naïve, primary tumor, or LN metastatic mice.

Comparison of late-stage LN metastatic cell lines to B16F0 cells revealed extensive differentially expressed genes. Annotation of genes involved in interferon responses (ISGs) revealed that a majority (64.6%) of the upregulated genes during LN metastasis are ISGs including key genes involved in antigen presentation (H2-K1), inflammatory responses (Isg15, Mx2), and immune regulation (Cd274) (Fig. 1B)^39^. Gene ontology (GO) analysis further revealed that many of the top ranked terms include interferon production and response terms and gene-set enrichment analysis (GSEA) highlighted that type I interferon responses as highly enriched in LN metastasis (Supp. Fig. 1D-G). To confirm the relevance of ISG expression in LN metastasis in humans, we analyzed the expression of genes involved in type I IFN response across primary tumor cells isolated from HNSCC patients. HNSCC patients with nodal metastases (N+) had higher expression of a majority of the type I IFN response genes compared with patients without nodal metastases (N0) (Fig. 1C).

The presence of type I and II interferon signatures in LN metastatic tumors across cancer types suggested a remodeling of the cytokine environment during dissemination. We next performed ex vivo analysis of type I interferons (IFNb and IFNa) and type II interferon (IFNy) in both the primary tumor site and in the tumor-draining LN. Both type I interferons were enriched in LN metastatic tumors compared to non-metastatic tumors and healthy skin tissue (Fig. 1D, E). In contrast, type II interferon was consistent across sites, resulting in a biased environment skewing towards type I interferons (Fig. 1F, G). Within metastatic LNs, type I interferon was enriched and type II interferon impaired, suggesting this biased cytokine environment is present at both the primary tumor and metastatic sites (Fig. 1H-K). Analysis of the HNSCC patient cohort, including RNA-seq FACS purified tumor cells and leukocytes, revealed that total interferon gene expression is enriched during LN metastasis with a bias towards leukocyte production (Fig. 1L, Supp. Fig. 1H-K).

Lymphocytes are potent interferon producers, particularly in response to recognition of cognate antigens^40^. To assess if anti-tumor T cell responses to tumors may be responsible for the ISG signature present in LN-metastatic tumors we performed a coculture of B16F0 cancer cells with polyclonal or OT-I CD8+ T cells, whose T cell receptor (TCR) specifically recognizes ovalbumin (OVA). H-2D^b^ and PD-L1 (surface markers for ISGs) were elevated on cancer cells only when OVA-expressing B16F0 tumor cells were co-cultured with OT-I T cells (Fig. 1M), suggesting lymphocyte recognition of tumors may induce this switch to type I interferon production. We next performed ex vivo cytokine analysis by flow cytometry for IFNa and IFNy to identify the cells responsible for cytokine secretion. Within metastatic tumors, we identified both CD4+ and CD8+ T cells that had increased release of IFNa, with the ratio of type I to type II interferon producing T cells showing a bias towards type I secretion (Fig. 1N-P). Within the tumor-draining LN, we found that lymphocytes broadly engaged in increased IFNa secretion including B cells and natural killer (NK) cells, while CD8+ T cells demonstrated impaired IFNy secretion (Figure 1Q-V). Together, these data suggest that lymphocyte-derived type I interferon is enriched in LN metastatic tumors and is responsible for interferon signatures across cancer types.

### Type I interferon promotes lymph node metastasis while impairing distant metastasis

Given the increase of type I IFN producing lymphocytes into LN-metastatic primary tumors and LN metastases and the potent induction of ISGs within LN-metastatic cancer cells, we sought to determine how type I IFNs contribute to metastasis. Metastasis occurs across long time scales, often months to many years in humans, suggesting phenotypes may arise during the course of tumorigenesis that enable spread throughout the body^41^. To address whether chronic exposure to type I IFNs from futile anti-tumor immune responses during tumor growth was sufficient to induce ISG signatures and drive metastasis, we evolved B16F0 and YUMM1.7 melanoma cell lines for four weeks in the presence of IFNa during culture before implanting the cells into mice to evaluate metastasis (Fig. 2A, Supp. Fig. 2A). In both the B16F0 and YUMM1.7 models, chronic exposure to IFNa was sufficient to impart a LN metastatic phenotype (Fig. 2B) without impacting primary tumor growth (Supp. Fig. 2B), driving both increased rates of metastasis and the development of larger metastases (Fig. 2C, D, Supp. Fig. 2C, D). Conditioning with type I IFNs during the equivalent time needed for tumor growth and metastasis in vivo was sufficient to induce LN metastasis, suggesting this cue provides the necessary program for colonization of LNs.

**Figure 2.**
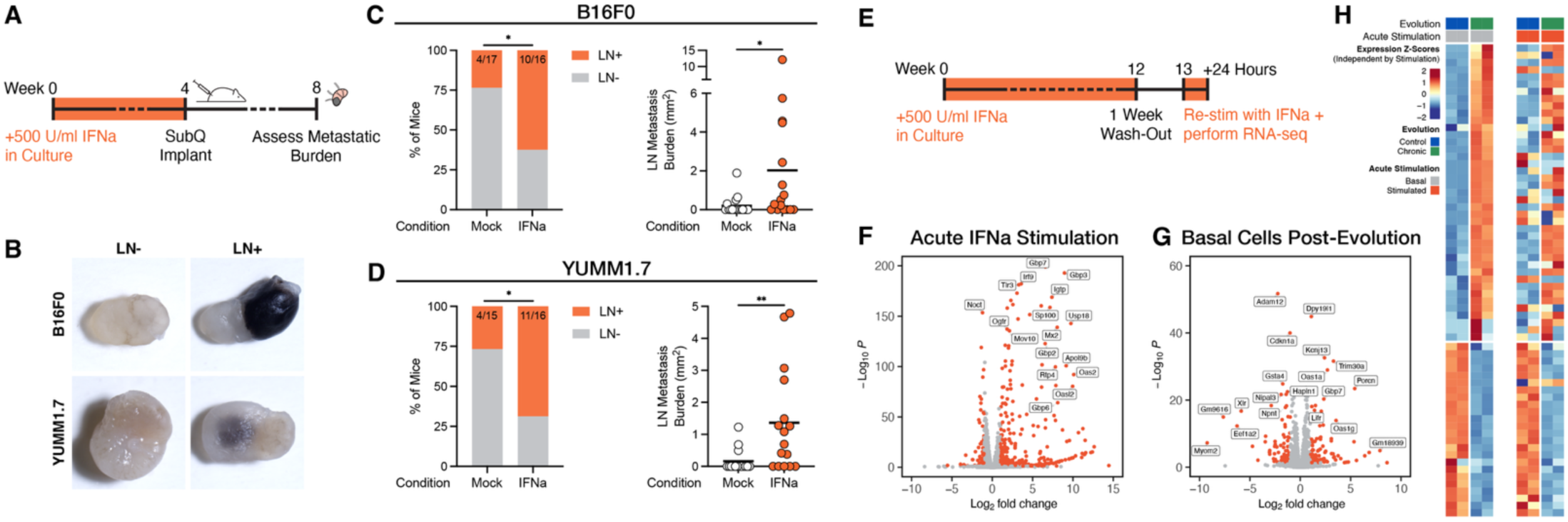
Chronic exposure to type I interferon is suXicient to drive LN metastasis. (A) Schematic of chronic interferon evolution and in vivo metastasis assessment. (B) Representative images of uninvolved and metastatic LNs from mice bearing B16F0 or YUMM1.7 tumors. (C) B16F0 and (D) YUMM1.7 frequency of LN metastases and LN metastasis burden according to mock or IFNa chronic evolution. (E) Schematic of extended chronic interferon evolution and wash-out for RNA-seq. (F) Volcano plot of differentially expressed genes for mock evolved cells receiving acute IFNa stimulation. (G) Volcano plot of differentially expressed genes comparing interferon to mock evolved cells without stimulation. (H) Heatmap of gene expression for differentially expressed genes from (G) for all conditions. Basal and stimulated groups are plotted on individual scales for visualization purposes.

To investigate the genetic mechanisms by which chronic type I interferon reprograms tumors, we performed transcriptional profiling. We exposed parental tumors to IFNa for a total of 12 weeks, before a week long wash-out period. Mock and conditioned cells were then re-stimulated with IFNa and subjected to bulk RNA-seq (Fig. 2E). Comparison of mock evolved cells that were acutely stimulated with IFNa revealed potent upregulation of many interferon-responsive genes (Fig. 2F). Comparison of chronic interferon evolved cells without stimulation to mock evolved cells revealed similar, albeit weakened, interferon transcriptional responses (Fig. 2G, H) with GO analysis indicating response to type I interferon as the top two terms (Supp. Fig. 2E). As we have previously demonstrated, ISGs enable evasion of NK cell killing and suppression of anti-tumor T cell responses, enabling cancer cells to colonize the LN^25^. Chronic type I interferon exposure contributes to a persistent transcriptional alteration maintaining ISG expression, likely through epigenetic remodeling resulting from persistent transcriptional activation at ISG loci^42,43^.

Given that type I interferon signaling was sufficient to drive LN metastasis, we next assessed if type I interferon was strictly necessary. The type I interferon receptor is composed of a heterodimer between Ifnar1 and Ifnar2 that forms upon binding of soluble type I interferons. We performed dual knockout of Ifnar1 and Ifnar2 in B16F0 or LN6-987AL cell lines, ablating the ability of the cells to transduce type I interferon signals (Supp. Fig. 3A-C). Elimination of type I interferon signaling did not significantly alter tumor growth rates (Fig. 3A). While tumor burden was consistent across genotypes, however, knockout of IFNAR1/2 in LN6-987AL cells reduced the number of LN metastases that formed and the LN metastatic burden in mice (Fig. 3B, C).

**Figure 3.**
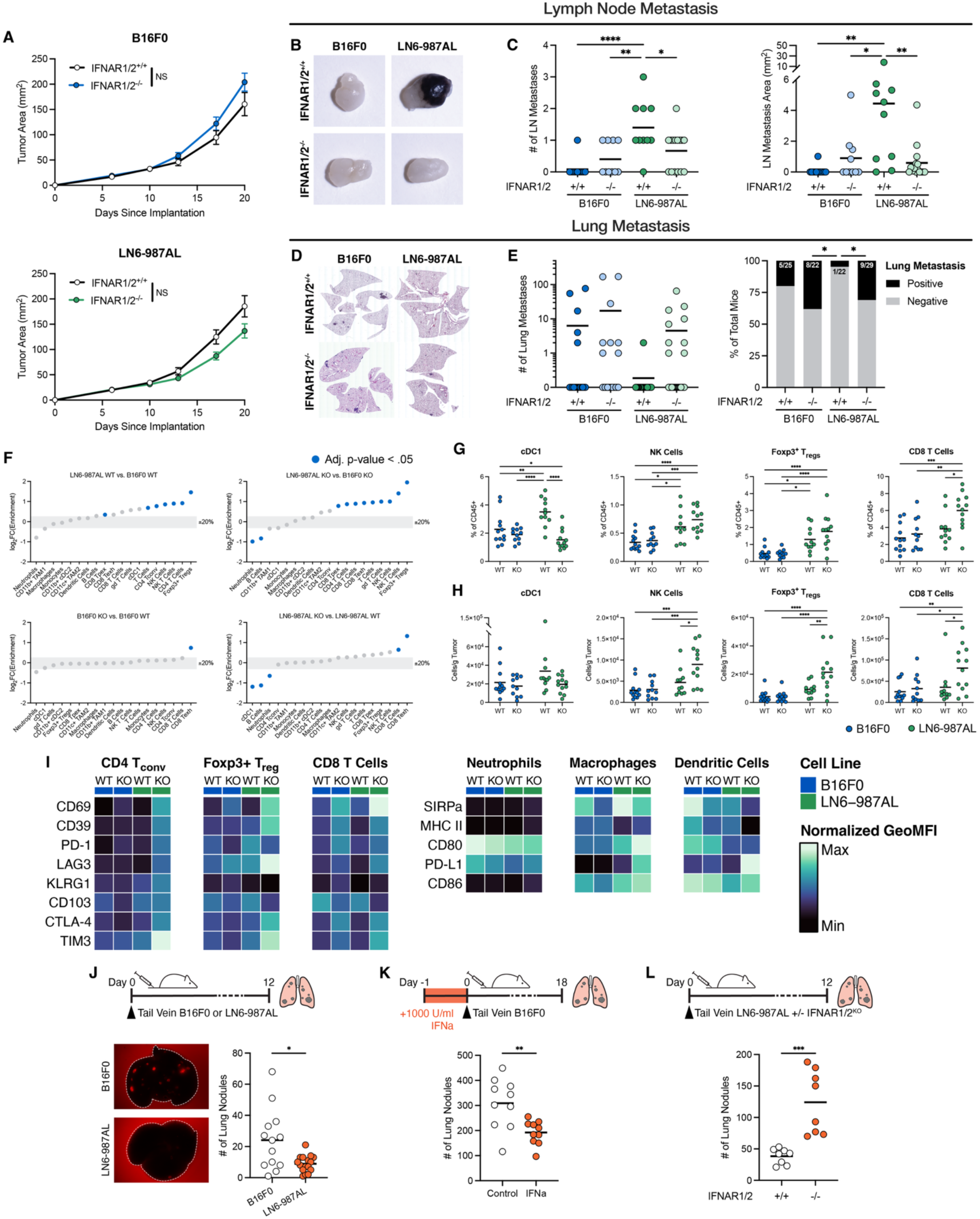
Type I interferon signaling has opposing roles in LN and distant organ metastasis. (A) Growth curves for B16F0 and LN6-987AL wildtype or IFNAR1/2 knockout tumors. (B) Representative LN images and (C) quantification of number of LN metastases and total burden. (D) Representative lung H&E images and (E) spontaneous lung metastasis counts and binary quantification. (F) Immune cell enrichment plots for B16F0 and LN6-987AL wildtype or IFNAR1/2 knockout tumors, with pairwise comparisons shown. Key immune cell frequencies (G) as percent of CD45+ cells in the tumor or (H) cell counts normalized to tumor mass. (I) Normalized geometric MFIs for functional immune markers on major immune cell types. (J) Representative lung images and lung metastasis counts comparing experimental metastasis of B16F0 or LN6-987AL cells. (K) Lung metastasis counts comparing experimental metastasis of basal or IFNa pre-treated B16F0 cells. (L) Lung metastasis counts comparing experimental metastasis of wildtype or IFNAR1/2 knockout LN6-987AL cells.

To determine whether the enhanced LN metastatic potential conferred by type I interferon signaling extends to metastasis to other tissues, we also assessed spontaneous lung metastasis in the same cohorts of mice bearing the IFNAR1/2 knockout LN6-987AL tumors. Unlike LN metastases, spontaneous lung metastases were exceedingly rare in mice bearing LN metastatic tumors. In contrast to its effects on reducing LN metastasis, however, ablation of interferon signaling rescued the lung metastatic potential of these tumors (Fig. 3D, E). Thus, type I interferon signaling is both necessary and sufficient to confer LN metastasis but opposes the formation of lung metastases.

To uncover the mediators of these divergent metastatic proclivities, we next explored what microenvironmental changes IFNAR1/2 knockout induced by performing flow cytometry profiling of the immune milieu of tumors (Supp. Fig. 4, 5). LN6-987AL tumors exhibited subtle changes in immune composition compared to parental B16F0 tumors, while knockout of IFNAR1/2 induced widespread remodeling of the immune microenvironment (Fig. 3F, Supp. Fig. 6, 7). cDC1 cells, capable of antigen cross-presentation and CD8 T cell priming^44^, are enriched within LN6-987AL tumors but are lost upon IFNAR1/2 knockout (Fig. 3G, H). There was also significant influx of lymphocyte populations including NK cells, Foxp3+ Tregs, and CD8 T cells within IFNAR1/2 knockout LN6-987AL tumors (Fig. 3G, H), suggesting alteration to immune surveillance may differentially affect cancer cells during metastasis. Similarly, functional marker expression on lymphocytes and myeloid cells was broadly altered upon IFNAR1/2 knockout (Supp. Fig. 6, 7), with loss of type I signaling in tumors contributing to enhanced expression of markers for antigen-experience and coinhibitory molecules (Fig. 3I).

Given the broad immune alterations that occur upon IFNAR1/2 knockout and the apparent opposing role for type I interferon in LN and spontaneous lung metastasis, we further dissected how interferon regulates distant metastasis through experimental tail vein challenges. Intravenous injection of LN6-987AL resulted in fewer lung nodules than B16F0, confirming the impaired lung metastatic potential of LN metastatic tumors observed before (Fig. 3J). Given that ISG expression is not the only modification that may be acquired during LN metastasis, we also asked whether pre-treatment of B16F0 cells to increase ISG expression alone was sufficient to impair lung metastasis, finding that solely increasing ISG expression was sufficient to phenocopy the impaired distant metastasis of LN metastatic cell lines (Fig. 3K). Finally, we asked whether lung metastasis could be rescued in LN metastatic cells by ablation of type I interferon signaling. Indeed, knockout of IFNAR1/2 in LN6-987AL cells drastically increased the number of lung nodules that formed (Fig. 3L). Collectively, these results demonstrate that type I interferon signaling imparts divergent metastatic phenotypes, simultaneously promoting LN metastasis while impairing metastatic seeding of distant sites. Tumor-intrinsic interferon signaling further appears to participate in a broad remodeling of the tumor microenvironment that is dependent on chronic signaling.

### Interferon signatures are selected for in LN metastases across cancers and enable immune evasion

In light of the apparent pro- and anti-metastatic activities of interferon signaling and ISG expression, we returned to our original in vivo selection approach for metastases, now extending it to look at simultaneous spread to LNs and lungs. We evolved paired metastases from the same tumor for B16F0 and performed two rounds of selection for LN and lung metastasis in the MOC2 oral squamous cell carcinoma model^45^ (Fig. 4A). This approach resulted in a triad cell line (primary tumor, LN metastasis, and lung metastasis) from B16F0 (Supp. Fig. 8A) and 18 LN and lung metastatic cell lines from MOC2 (Supp. Fig. 8B). Surface markers for ISG expression were increased on LN-derived and decreased on lung-derived B16F0 metastases compared to their matched primary tumor (Fig. 4B), supporting the notion that ISG expression is selected for and against during cell dissemination from the primary tumor site. NK cell killing has previously been established as a substantial barrier to survival of circulating cells during hematogenous^46^ and lymphatic metastatic spread^25^ and is regulated in part by interactions between NK inhibitory receptors and MHC I, a key ISG. Consistent with altered expression of the classical MHC-I molecule H-2D^b^ during metastasis, we find that NK cell lysis of LN-derived cell lines is impaired and lung-derived cell lines is enhanced (Fig. 4C). These same ISG expression patterns and NK cell cytotoxicity susceptibilities are exhibited in our MOC2 metastasis model (Fig. 4D, E Supp. Fig. 8C, D), demonstrating the generalizability of these findings across tumor types. Supporting the notation that MHC I is contributing to altered NK cell killing, MHC-I expression negatively correlates with NK cell killing across LN and lung-derived MOC2 metastatic cell lines (Fig. 4F).

**Figure 4.**
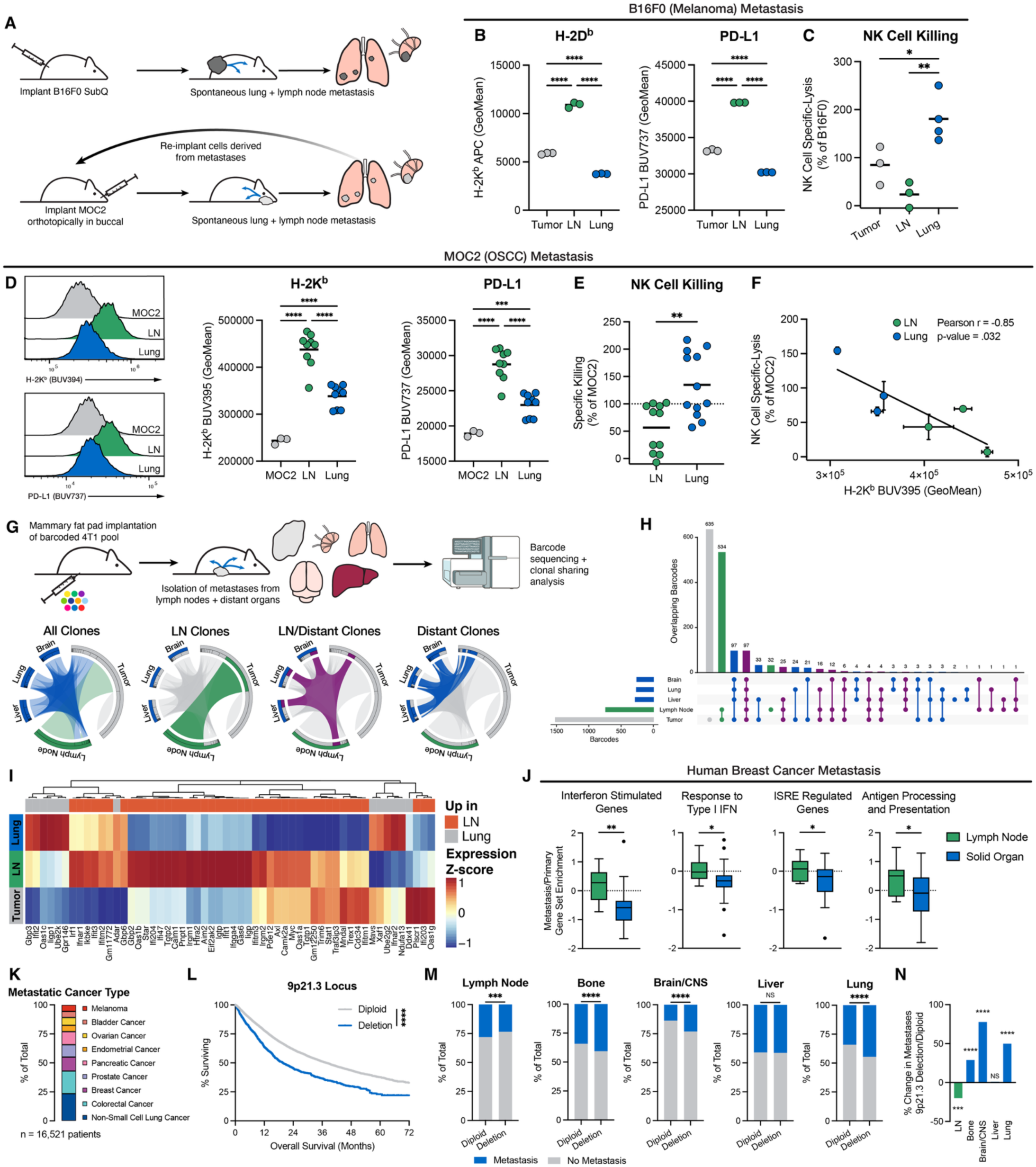
Divergent ISG expression between LN and distant organ metastasis accounts for independent clonal seeding. (A) Schematic of metastatic evolution approach for triad B16F0 cell lines and second generation MOC2 cell lines. (B) H-2Kb and PD-L1 expression on B16F0 triad cell lines and (C) NK cell killing efficiency. (D) H-2Kb and PD-L1 expression on second generation MOC2 metastatic cell lines and (E) normalized NK cell killing efficiency. (F) Correlation of NK cell killing efficiency with H-2Kb expression on MOC2 metastatic cell lines. (G) Overview of 4T1 metastatic barcoding approach and clonal overlap between primary tumor, LN, and distant organ sites. (H) Upset plot quantifying barcode overlap between sites in (G). (I) Type I interferon responsive gene expression for B16F0 triad cell lines. (J) Interferon-related gene set enrichment scores for human breast cancer paired primary tumor and LN or solid organ metastases. (K) Overview of MSK-MET cohort subset analyzed. (L) Survival curve for MSK-MET cohort stratified by predicted 9p21.3 diploid or deletion. (M) Frequency of metastasis to various sites stratified by predicted 9p21.3 deletion and (N) percent change in site frequency between deletion or diploid subgroups.

The divergent ISG profiles and effects of IFNAR signaling of LN and lung metastases are suggestive of distinct clonal selection biases in LNs and distant tissues, consistent with clinical evidence of independent clonal origins between metastases in these tissues. Thus, we further explored the clonal basis of metastatic dissemination. We leveraged a dataset of 4T1 breast cancer cells that were transduced with a library of unique DNA barcodes prior to implantation in mice followed by barcode sequencing of metastases^13^ (Fig. 4G). Barcode overlap analysis between tissue sites revealed that while numerous tumor clones metastasized to only LNs, clones that formed distant metastases without detectable LN metastases were as abundant as those that metastasized to both LNs and distant tissues (Fig. 4H). This finding is consistent with reports in humans suggesting that LNs receive the most unique metastatic cells and that distant metastatic sites are frequently seeded by clones that never seed the LN^11,20^. ISG expression modulation according to the tissue site of metastasis is thus a likely basis for the routine independent seeding of metastases from primary sites in cancer patients.

To further investigate this concept, we evaluated broader ISG profiles in our triad family of cell lines derived from B16F0 by bulk RNA-seq. We again found that in metastases derived from the same primary tumor, interferon I responsive gene expression was increased in cells in the LN and decreased in those from the lung (Fig. 4I). We expanded this analysis to rare paired primary tumor and metastatic samples from breast cancer patients that underwent bulk RNA-seq, evaluating the change in metagene expression scores from the primary to distinct metastatic sites. Consistent with our data in murine models, LN metastasis samples exhibited an increase in metagene expression scores for processes like response to type I interferon and antigen presentation, while distant organ metastases exhibited a decrease, (Fig. 4J).

Due to the limited gene expression data available for metastatic cancer patients, we also looked at genetic sequencing data, enabling us to profile the sites of metastasis for 16,521 patients across 9 major cancer types with metastatic disease from the MSK-MET project^47^, assayed on the MSK-IMPACT targeted sequencing platform^48^ (Fig. 4K). Since ISG expression data was not available, we instead looked at the occurrence of deletions at the 9p21.3 locus (predicted by CDKN2A loss by MSK-IMPACT sequencing), which contains many of the type I interferon genes and is associated with impaired immune infiltration to tumors^49,50^. Subsetting only on patients with metastases, deletion of 9p21.3 was associated with a reduction of 18 months median survival (48% reduction) across all cancer types (Fig. 4L). Analysis of the site of metastasis for patients stratified by diploid genotype or loss of 9p21.3 revealed that patients with deletions had fewer LN metastases but increased metastases in the bone, brain, and lungs (Fig. 4M).

Unlike the pro-metastatic effect of 9p21.3 deletion on distant metastatic sites, loss of this locus (including a range of interferon genes) starkly reduced LN metastasis, in alignment with our findings that type I interferons act to drive LN metastasis (Fig. 4N), adding nuance to the solely pro-metastatic roles previously identified^51^. Together, these findings suggest that high ISG expression is a trait that is modulated during metastasis to LNs or other distant sites, even from the same primary tumor. Thus, we demonstrate that ISG expression accounts for the divergent seeding pattern of tumors, offering a mechanistic rationale for why LN and distant metastases are often derived from independent clones.

Primary tumor clones that receive interferon stimulation become destined for LNs while those that avoid stimulation become suitable for seeding of distant organs, with the adaptations for the LN making them poor at seeding additional downstream sites.

### Lymph node metastasis and interferon signaling sensitize tumors to immune checkpoint blockade

ICB therapy has yielded immense success for many cancers in the clinical setting^52–54^, but resistance remains a significant challenge for the majority of patients with distinguishing clinical factors during disease progression poorly understood^55^. LN metastases can promote the generation of Tregs that contribute to tolerance to the tumor and induce unproductive lymphocyte states during immune checkpoint blockade (ICB)^25–27^. Direct functional testing for the impact of LN metastases on ICB responses beyond associations remains sparse. Thus, we first tested if LN metastasis altered the impact of ICB therapy on primary tumor growth, using a regimen targeting both CTLA-4 and PD-L1 (Fig. 5A). Much like the majority of cancer patients, the parental B16F0 model is refractory to ICB therapy (Fig. 5B)^55^. In contrast, LN metastatic tumors exhibited enhanced response to ICB therapy, demonstrating significantly delayed tumor growth, resulting in a two-fold in tumor size (Fig. 5C), and a similar response is achieved with anti-PD-L1 monotherapy (Supp. Fig. 9A, B). Though correlative data in human cohorts have previously suggested LN metastasis weakened responses to ICB through decreases of progenitor exhausted and intermediate exhausted CD8 T cells^27^, we find that ISG signatures acquired during LN metastases render patients more susceptible to ICB therapy, likely due to ISG-driven expression of targetable checkpoints like PD-L1 and heightened presentation of antigen directly at sites of T cell priming through MHC I expression.

**Figure 5.**
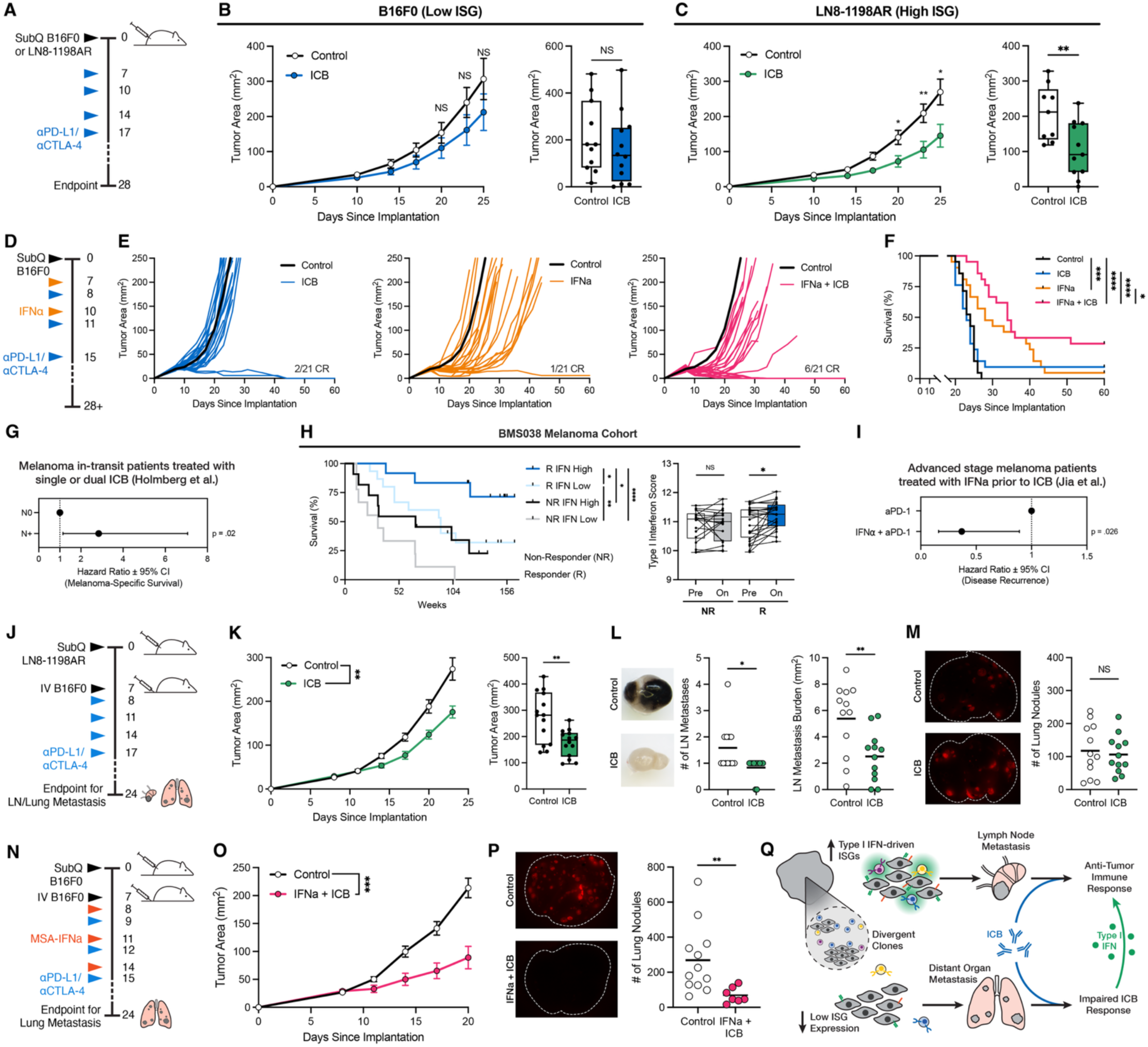
LN metastasis induces vulnerability to ICB therapy and can be mimicked through type I interferon delivery. (A) Schematic of ICB dosing schedule. (B) B16F0 and (C) LN8-1198AR tumor growth curves and individual tumor volumes at Day 23. (D) Schematic of intratumoral IFNa sensitization and ICB combination therapy dosing schedule. (E) Individual tumor growth curves for each mouse compared to mean of control tumors. (F) Survival curves for mice in (E). (G) Hazard-ratio plot for melanoma-specific survival for patients treated with ICB according to nodal involvement (n = 287 patients). (H) BMS038 melanoma cohort treated with ICB with pre/post-treatment bulk RNA-seq. Left, survival plot stratified by responder/non-responder status and pre-therapy interferon signaling score. Right, type I interferon signaling score according to responder/non-responder status grouped by pre/on-therapy. (I) Hazard-ratio plot for disease recurrence for advanced stage melanoma patients treated with aPD-1 therapy, stratified by previous treatment with IFNa (n = 56 patients). (J) Schematic of ICB dosing and tail vein injection schedule to simultaneously evaluate LN and lung metastasis in LN8-1198AR model. (K) Tumor growth curves and end-point tumor area, (L) Representative LN metastasis images and quantification of LN metastasis burden, and (M) representative lung metastasis images and quantification of burden for mice in (J). (N) Schematic of systemic extended half-life IFNa and ICB dosing for lung metastasis evaluation. (O) Tumor growth curves and (P) representative lung metastasis images and quantification of burden for mice in (N). (Q) Diagram overview of proposed mechanism, including individual clones from primary tumors receiving differential immune attack, inducing high ISG expression patterns in some cells that seed LN metastases and low ISG expression in other that seed distant organ metastases, giving rise to the divergent seeding and clonal pattern of cancer metastasis. Due to high ISG expression, only LN metastases but not distant metastases are capable of responding to ICB therapy, but delivery of type I interferon is sufficient to convert resistant metastatic lesions to ICB-susceptible tumors, enabling clearance.

Based on our evidence suggesting that the interferon signatures acquired during LN metastasis contribute to ICB response, we asked whether nonresponsive primary tumors can be sensitized to ICB through type I interferon exposure. We administered intratumoral type I interferon to mice bearing B16F0 parental tumors treated with dual ICB (Fig. 5D, Supp. Fig. 9C, D)^56^. Here, we found that again ICB had limited activity on B16F0 tumors while delivery of IFNa elicited a more potent, yet brief response (Fig. 5E). It was only in the case when IFNa and ICB were delivered in combination with each other that tumor growth was delayed and 29% of mice achieved complete responses (Fig. 5E, F), resulting in significantly improved survival no observable toxicity (Supp. Fig. 9E, F). In light of this improvement in survival and our discovery that LN metastases exhibit higher levels of type I interferon signaling, we interrogated a series of human cohorts and found evidence that LN metastasis enhanced outcomes during ICB therapy. A multicenter retrospective analysis of stage III in-transit melanoma patients suggests that LN metastasis was associated with enhanced melanoma-specific survival (Fig. 5G)^57^. To directly evaluate relationships between interferon signaling and checkpoint response, we predicted type I interferon responsiveness for patients in the BMS038 melanoma cohort^58^ using their pre-ICB therapy samples, allowing us to stratify survival for non-responders and responders according to interferon signaling (Fig. 5H). We also computed a type I interferon response metagene score for patient samples pre- and on-therapy, finding that only responders to ICB therapy exhibited enhanced scores during therapy administration. Finally, in a cohort of late-stage melanoma patients, it was found that prior exposure to pegylated IFNa therapy significantly decreased the risk of disease recurrence for patients (Fig. 5I)^59^.

While the association between LN metastasis and improved primary tumor responses to ICB therapy was unexpected, metastasis, not primary tumor burden, results in the majority of cancer deaths^1,2^. Thus, we sought to determine whether type I interferon could be harnessed for the treatment of ICB-resistant metastatic disease. We first assessed the ability of ICB therapy to reduce metastatic burden at both LN and lung sites through simultaneous spontaneous LN metastasis and experimental lung metastasis assays (Fig. 5J). As we found before, LN metastatic primary tumors were responsive to dual ICB therapy (Fig. 5K). Similarly, LN metastasis was effectively targeted by ICB, resulting in fewer metastases and reduced metastatic burden (Fig. 5L). As has been demonstrated previously for many tumor types, distant metastases in the lung were non-responsive to ICB therapy^55^(Fig. 5M), suggesting alternative strategies must be used to overcome the burden of metastasis. Building upon our previous efforts to sensitize primary tumors to ICB by delivery of IFNa to enforce an ISG-high LN metastasis-like state, we designed a half-life extended IFNa molecule for systemic delivery through fusion with mouse serum albumin (MSA)^60^ (Supp. Fig. 9G, H). Sequential systemic delivery of extended half-life IFNa and ICB with experimental lung metastasis (Fig. 5N) was capable of impairing primary tumor growth without toxicity as we demonstrated through intratumoral delivery earlier (Fig. 5O, Supp. Fig. 9I, J). Systemic delivery of combination therapy was also sufficient to enable clearance of previously established lung metastases, significantly reducing metastatic burden unlike ICB alone (Fig. 5P). Thus, by harnessing the intrinsic ISG-dependence of naturally ICB-responsive LN metastases, type I interferon is capable of converting distant metastases from immunotherapy-refractory to immunotherapy-responsive (Fig. 5Q).

## Discussion

Distant organ metastases are frequently composed of tumor clones that appear absent from LN metastases in mouse models and humans^10,11,13,61,62^. This clonal exclusivity stands in contrast to previous theories suggesting that cancer cells needed to colonize LNs in order to acquire additional traits that aid further dissemination^6^. These observations of independent seeding have been especially perplexing given the greater diversity of colonizing tumor clones in the LN, likely due to a high rate of cellular drainage^20^.

Nonetheless, LNs and distant tissues, and lymph and blood themselves, pose disparate microenvironmental barriers that can sculpt the phenotypes of metastases^63^. Here, we discover that not only do distinct molecular mechanisms promote LN metastasis, but that these mechanisms are detrimental to seeding distant sites. Tumor-intrinsic ISG programs induced by type I interferon derived from the immune system enable colonization of LNs through evasion and suppression programs such as expression of MHC I, which inhibits NK cell cytotoxicity. These same ISG programs directly impair the ability of tumor cells to form overt distant metastases. This ISG sensitivity is consistent with previous work that found silencing of ISG programs was a crucial step in the development of bone metastases and mediated evasion of immune surveillance^64,65^. Low ISG expression in tumor cells likely aids survival at distant sites through a variety of pathways, including minimizing tumor-immune interactions with myeloid cells, CD8 T cell escape through loss of MHC I, and downregulation of NK-activating ligands^66–68^. Thus, type I interferon-induced ISG programs in tumor cells have opposing roles in the formation of LN or distant organ metastasis, accounting for the comparatively high-rate of independent seeding of LN or distant metastases in mouse models and human patients.

While previous work in both mouse models and clinical cohorts has suggested that de-escalation of traditionally aggressive lymphadenectomy in cancer patients best preserves tumor-draining LNs that are critical for mounting a systemic response to immune checkpoint blockade, immune correlate markers have suggested this approach may be detrimental once LN metastases have formed^24,27,28^. Here, we demonstrate that though markers of dysfunctional lymphocyte responses are enhanced within the tumor upon LN metastasis, dissemination of cancer to LNs can result in enhanced responses to ICB therapy.

We suggest that this is due to a combination of factors, including the acquisition of ISG programs in tumor cells during the process of forming LN metastases such as MHC I and PD-L1 expression and the close interaction between cancer and immune cells in the LN, enabling effective lymphocyte responses to be generated. The discordance between immune correlate markers of lymphocyte dysfunction and productive responses to therapy may be due to the overlapping nature of these markers as both signals of immune activation and dysfunction^69,70^. Additionally, there may be limitations to the degree to which an involved LN is effaced with tumor that still preserves the requisite architecture for mounting immunity. Thus, this work extends the argument for LN-sparing surgical approaches for early-stage cancer patients receiving ICB therapy to those with regional LN metastases, though further work dissecting ICB responses among patients with nodal metastases prior to further dissemination is needed.

Once thought to be uniformly associated with productive anti-tumor responses, interferon signaling is now appreciated to play a more nuanced role in tumor regression and response to treatment. In the context of radiotherapy, type I interferon can be induced through cancer-intrinsic sensing of damage signals, promoting myeloid cell antigen presentation and the generation of effective anti-tumor lymphocyte responses^71,72^. In the context of de novo ICB-resistance, interferons can act in a similar fashion, eliciting productive immune responses. However, in the context of acquired ICB-resistance, chronic interferons play a negative role, contributing to T cell exhaustion which can be targeted successfully through JAK1 inhibition, preventing chronic interferon signaling^42,73,74^. Indeed, a recent study evaluating humoral correlates to ICB response uncovered that, unlike the infectious disease context, patients who produce anti-type I interferon auto-antibodies are among the strongest responders to ICB^75^. Thus, these divergent roles of type I interferon afford opportunities for therapeutically modulating ICB responses in otherwise unresponsive patient populations.

We demonstrate that though LN metastases and their originating primary tumors are more susceptible to ICB therapy, metastases to other sites remain resistant to ICB, likely due in part to downregulation of ISG programs. Delivery of type I interferon enabled sensitization of metastases to ICB therapy, converting once resistant tumors to immunotherapy-sensitive targets. A key limitation to this approach is the relatively high toxicity profile of previous type I interferon molecules in humans^76^, however, previous work on the development of biased- and synthetic-cytokines has demonstrated that toxicity of endogenous cytokines for cancer therapy need not be accepted as immutable and may yield effective molecules for sensitizing metastases to ICB therapy^77,78^. Uncovering the basis for divergent metastatic seeding and the benefit to LN metastasis during ICB therapy enabled us to design a proof-of-concept approach for sensitizing distant metastases to ICB therapy by converting them to LN-like metastases with type I interferon. Thus, the LN-specific type I interferon metastasis axis represents an archetype for how the defining drivers of clonal architecture can be exploited to improve treatments for stage IV metastatic disease.

## Methods

### Key Resources Table

| REAGENT or RESOURCE | SOURCE | IDENTIFIER |
| --- | --- | --- |
| Antibodies |  |  |
| anti-mouse CTLA-4 (Clone 9H10) | BioXCell | Cat# BE0131<br>RRID:AB_10950184 |
| anti-mouse PD-L1 (Clone 10F.9G2) | BioXCell | Cat# BE0101<br>RRID:AB_10949073 |
| anti-mouse CD16/32 (Clone 2.4G2) | BioXCell | Cat# BE0307<br>RRID:AB_2736987 |
| anti-human/mouse Phospho-Stat1 (Tyr701) Alexa Fluor 647 (Clone 58D6) | Cell Signaling Technology | Cat# 8009S<br>RRID:AB_10860764 |
| anti-mouse H-2Db APC (Clone KH95) | Biolegend | Cat# 111514 |
| anti-mouse H-2Kb APC (Clone AF6-88.5) | Biolegend | Cat# 116518 |
| anti-mouse H-2Kb BUV395 (Clone AF6-88.5) | BD Biosciences | Cat# 742864 |
| anti-mouse H-2Kb PE (Clone AF6-88.5) | Biolegend | Cat# 116508 |
| anti-mouse PD-L1 BUV737 (Clone 10F.9G2) | BD Biosciences | Cat# 568826 |
| anti-mouse B220 BV650 (Clone RA3-6B2) | Biolegend | Cat# 103241 |
| anti-mouse B220 PE-Fire 810 (Clone RA3-6B2) | Biolegend | Cat# 103287 |
| anti-mouse CD103 PE-Dazzle 594 (Clone 2E7) | Biolegend | Cat# 121430 |
| anti-mouse CD11b FITC (Clone M1/70) | Biolegend | Cat# 101206 |
| anti-mouse CD11c FITC (Clone N418) | Biolegend | Cat# 117306 |
| anti-mouse CD11c PE (Clone N418) | Biolegend | Cat# 117308 |
| anti-mouse CD25 BUV395 (Clone PC61) | BD Biosciences | Cat# 564022 |
| anti-mouse CD39 BV510 (Clone Y23-1185) | BD Biosciences | Cat# 567265 |
| anti-mouse CD4 BUV805 (Clone RM4-5) | Thermo Fisher Scientific | Cat# 368-0042-82 |
| anti-mouse CD44 BV785 (Clone IM7) | Biolegend | Cat# 103059 |
| anti-mouse CD45 BUV496 (Clone 30-F11) | BD Biosciences | Cat# 569673 |
| anti-mouse CD62L APC-R700 (Clone MEL-14) | BD Biosciences | Cat# 565159 |
| anti-mouse CD69 BUV737 (Clone H1.2F3) | BD Biosciences | Cat# 612793 |
| anti-mouse CD80 BV650 (Clone 16-10A1) | Biolegend | Cat# 104732 |
| anti-mouse CD86 BUV395 (Clone GL1) | BD Biosciences | Cat# 564199 |
| anti-mouse CD8a BUV615 (Clone 53-6.7) | BD Biosciences | Cat# 613004 |
| anti-mouse CD8a Spark Blue 574 (Clone 53-6.7) | Biolegend | Cat# 100794 |
| anti-mouse CTLA-4 PE-Cy7 (Clone UC10-4B9) | Biolegend | Cat# 106314 |
| anti-mouse F4/80 BV421 (Clone BM8) | Biolegend | Cat# 123131 |
| anti-mouse Foxp3 BV421 (Clone FJK-16S) | Thermo Fisher Scientific | Cat# 404-5773-82 |
| anti-mouse gdTCR PE-Cy5 (Clone eBioGL3) | Thermo Fisher Scientific | Cat# 15-5711-82 |
| anti-mouse IFNα FITC (Clone RMMA-1) | PBL Assay Science | Cat# 22100 |
| anti-mouse IFNγ APC (Clone XMG1.2) | Biolegend | Cat# 505810 |
| anti-mouse KLRG1 PE (Clone 2F1) | Thermo Fisher Scientific | Cat# 12-5893-82 |
| anti-mouse LAG3 PerCP-Cy5.5 (Clone C9B7W) | Biolegend | Cat# 125212 |
| anti-mouse Ly6C R718 (Clone AL-21) | BD Biosciences | Cat# 566987 |
| anti-mouse Ly6G PE-Cy7 (Clone 1A8) | Biolegend | Cat# 127617 |
| anti-mouse MHC II (I-A/I-E) RB705 (Clone M5/114.15.2) | BD Biosciences | Cat# 570619 |
| anti-mouse NK1.1 PE-Cy7 (Clone PK136) | Biolegend | Cat# 108714 |
| anti-mouse NK1.1 BUV615 (Clone PK136) | Thermo Fisher Scientific | Cat# 366-5941-82 |
| anti-mouse PD-1 BV421 (Clone 29F.1A12) | Biolegend | Cat# 135217 |
| anti-mouse PD-1 BV605 (Clone 29F.1A12) | Biolegend | Cat# 135219 |
| anti-mouse PD-L1 BUV737 (Clone 10F.9G2) | BD Biosciences | Cat# 568826 |
| anti-mouse SIRPα AF647 (Clone P84) | Biolegend | Cat# 144028 |
| anti-mouse TCF7 AF647 (Clone S33-966) | BD Biosciences | Cat# 566693 |
| anti-mouse TCRβ RB705 (Clone H57-597) | BD Biosciences | Cat# 570604 |
| anti-mouse TCRβ BV711 (Clone H57-597) | Biolegend | Cat# 109243 |
| anti-mouse TIM3 APC-Fire 750 (Clone RMT3-23) | Biolegend | Cat# 119738 |
| anti-mouse XCR1 BV785 (Clone ZET) | Biolegend | Cat# 148225 |
| Bacterial and Virus Strains |  |  |
| Stable Competent *E. coli* | New England Biolabs | Cat# C3040H |
| Biological Samples |  |  |
| Chemicals, Peptides, and Recombinant Proteins |  |  |
| Universal Type I IFN (Hybrid Human IFNα) | PBL Assay Science | Cat# 11200 |
| Universal Type I IFN (IFN-αA/D(BgIII)) | Sperber et al.<sup>56</sup> and this paper | Supplementary Table 5 |
| C/N-term MSA-Universal Type I IFN (IFN-αA/D(BgIII)) | This paper | Supplementary Table 5 |
| Hygromycin B | Thermo Fisher Scientific | Cat# 10687010 |
| Lipofectamine 3000 | Thermo Fisher Scientific | Cat# L3000015 |
| 16% Paraformaldehyde Solution | Electron Microscopy Sciences | Cat# 15710 |
| Expi293 Expression Medium | Thermo Fisher Scientific | Cat# A1435101 |
| Brilliant Stain Buffer | BD Biosciences | Cat# 566349 |
| LIVE/DEAD Fixable Blue Dead Cell Stain | Thermo Fisher Scientific | Cat# L34962 |
| Collagenase 4 | Worthington | Cat# LS004189 |
| DNase I | Sigma-Aldrich | Cat# DN25-1G |
| PMA/Ionomycin Cell Activation Cocktail (without Brefeldin A) | BioLegend | Cat# 423301 |
| PMA/Ionomycin Cell Activation Cocktail (with Brefeldin A) | BioLegend | Cat# 423303 |
| Monensin Solution | BioLegend | Cat# 420701 |
| Calcein AM | Thermo Fisher Scientific | Cat# C1430 |
| AccuCount Fluorescent Particles (5-5.9 um) | Spherotech | Cat# ACFP-50-5 |
| Trypsin-EDTA (.25%) with phenol red | Thermo Fisher Scientific | Cat# 25200114 |
| TrypLE Express Enzyme | Thermo Fisher Scientific | Cat# 12605010 |
| Accutase | Innovative Cell Technologies | Cat# AT104 |
| High-glucose DMEM with phenol red, glutamine-free | Thermo Fisher Scientific | Cat# 11960069 |
| RPMI 1640 with glutamine, HEPES, and phenol red | Thermo Fisher Scientific | Cat# 22400105 |
| High-glucose DMEM, glutamine/phenol red-free | Thermo Fisher Scientific | Cat# 31053028 |
| Fetal Bovine Serum | R&D Systems | Cat# S11150 |
| 10% Neutral Buffered Formalin Solution | Sigma-Aldrich | Cat# HT501128-4L |
| Isoflurane | Dechra | — |
| 16% Aqueous Paraformaldehyde | Electron Microscopy Sciences | Cat# 15710 |
| HEPES | Thermo Fisher Scientific | Cat# 15630080 |
| Penicillin-Streptomycin | Thermo Fisher Scientific | Cat# 15140163 |
| L-Glutamine | Thermo Fisher Scientific | Cat# 25030164 |
| MEM Non-Essential Amino Acids Solution | Thermo Fisher Scientific | Cat# 11140050 |
| Beta-Mercaptoethanol | Thermo Fisher Scientific | Cat# 31350010 |
| Critical Commercial Assays |  |  |
| Mouse IFNα All Subtype High-Sensitivity ELISA Kit | PBL Assay Science | Cat# 42115 |
| Mouse IFNβ High-Sensitivity ELISA Kit | PBL Assay Science | Cat# 42410 |
| LEGEND MAX Mouse IFNγ ELISA Kit | BioLegend | Cat# 430807 |
| ExpiFectamine 293 Transfection Kit | Thermo Fisher Scientific | Cat# A14525 |
| Superdex 75 Increase column 10/300 GL | Cytiva | Cat# 29148721 |
| EasySep Mouse NK Cell Isolation Kit | StemCell Technologies | Cat# 19855 |
| EasySep Mouse CD8+ T Cell Isolation Kit | StemCell Technologies | Cat# 19853 |
| eBioscience Foxp3 Fixation/Permeabilization Kit | Thermo Fisher Scientific | Cat# 00-5523-00 |
| Cytofix/Cytoperm Fixation/Permeabilization Kit | BD Biosciences | Cat# 554714 |
| Deposited Data |  |  |
| Bulk RNA LN Lines | Reticker-Flynn et al.<sup>25</sup> | GEO: GSE117529 |
| HNSCC Bulk RNA | Reticker-Flynn et al.<sup>25</sup> and Brennan et al.<sup>79</sup> | GEO: GSE113839 |
| B16F0-868 Bulk RNA | This paper |  |
| B16F0/YUMM1.7 Chronic IFN Bulk RNA | This paper |  |
| AURORA BIG RNA | Garcia-Recio et al.<sup>80</sup> | GEO: GSE209998 |
| BMS038 RNA-seq | Riaz et al.<sup>58</sup> | GEO: GSE91061 |
| Experimental Models: Cell Lines |  |  |
| B16F0 | ATCC | Cat# CRL-6322<br>RRID:CVCL_0604 |
| B16F0-cOVA | Reticker-Flynn et al.<sup>25</sup> | — |
| B16F0-tdTomato | Reticker-Flynn et al.<sup>25</sup> | — |
| NBF0LN6-987AL | Reticker-Flynn et al.<sup>25</sup> | — |
| NBF0LN6-987AL-tdTomato | Reticker-Flynn et al.<sup>25</sup> | — |
| NBF0LN8-1198AR | Reticker-Flynn et al.<sup>25</sup> | — |
| MOC2 | Laboratory of John Sunwoo | RRID:CVCL_ZD33 |
| YUMM1.7 | Laboratory of Amanda Kirane | RRID:CVCL_JK16 |
| Expi293F | Thermo Fisher Scientific | Cat# A14527 |
| B16F0-868Tumor | This paper | — |
| B16F0-868LN | This paper | — |
| B16F0-868Lung | This paper | — |
| MOC2 LN Lines | This paper | — |
| MOC2 Lung Lines | This paper | — |
| Experimental Models: Organisms/Strains |  |  |
| C57BL/6NCrl | Charles River Laboratories | RRID:IMSR_CRL:027 |
| C57BL/6-Tg(TcraTcrb)1100Mjb/J | Jackson Laboratories | RRID:IMSR_JAX:003831 |
| Oligonucleotides |  |  |
| Cas9 sgRNAs for PX458 | This paper | Supplementary Table 1 |
| Recombinant DNA |  |  |
| pSpCas9(BB)-2A-GFP (PX458) | Addgene, Ran et al.<sup>81</sup> | Cat# 48138 |
| pUltra-Chili-Luc | Addgene, Fujiwara et al.<sup>82</sup> | Cat# 48688 |
| psPAX2 | Addgene | Cat# 12260 |
| pCMV-VSV-G | Addgene, Stewart et al.<sup>83</sup> | Cat# 8454 |
| pD649 | ATUM | Cat# PD649 |
| Software and Algorithms |  |  |
| FlowJo (Version 10.10.0) | Becton, Dickinson & Company | — |
| Prism (Version 10.5.0) | GraphPad | — |
| Other |  |  |
| NovoCyte Quanteon | Agilent | — |
| NovoCyte Penteon | Agilent | — |
| Cytoflex-S | Beckman Coulter | — |
| 5-Laser Aurora | Cytek Biosciences | — |
| FACSAria II | BD Biosciences | — |
| EVOS M7000 | Thermo Fisher Scientific | — |
| SMZ1270i Stereo Dissection Microscope | Nikon | — |
| Countess 3 | Thermo Fisher Scientific | — |
| Spark Plate-Reader | Tecan | — |

### Animal Care

All animal studies were reviewed and approved by the Stanford University Institutional Animal Care and Use Committee under protocol APLAC-34497. C57BL/6 mice were purchased from Charles River Laboratories (C57BL/6NCrl, Stock #027) and allowed to acclimate to their housing conditions for 3-7 days prior to beginning experiments. OT-I mice (C57BL/6-Tg(TcraTcrb)1100Mjb/J) were purchased from Jackson Laboratories and bred for experiments. Animals were housed in specific pathogen-free conditions with a 12-hour light/dark cycle and approximately 50% humidity and 21 C temperature. Cages were changed every two weeks and animals were provided with ad libitum access to chow and non-acidified water. Animal experiments were initiated when mice were 7-9 weeks of age and experimental groups within experiments were co-housed. Either females or males were used in experiments as indicated to match the sex of the tumor cell line used.

### Cell Line Culture

B16F0 (#CRL-6322) cells were obtained from ATCC, YUMM1.7 cells were a gift from Amanda Kirane, and MOC2 cells were a gift from John Sunwoo. LN metastatic cell lines derived from B16F0 were generated as previously described^25,84^ or as described below for 868T, 868LN, and 868L lines. B16F0 and derivative lines, YUMM1.7, and MOC2 and derivative lines were grown in High-Glucose DMEM supplemented with 10% FBS, 4 mM L-Glutamine, and 100 U/ml Penicillin-Streptomycin (cDMEM). All cell lines were routinely tested for mycoplasma contamination by PCR and confirmed negative.

### In vivo Tumor models

B16F0 and LN metastatic cell lines were grown in mice as previously described and is briefly described here^84^. YUMM1.7 cells were grown and implanted identically to B16F0 cell lines. B16F0 and LN metastatic cell lines were always implanted in female mice and YUMM1.7 cells always implanted in male mice. Cells were grown for a minimum of two passages after thawing and detached with Accutase (Innovative Cell Technologies), quenched with cDMEM, and resuspended in phenol red-free DMEM. Cell suspensions were counted twice using a Countess 3 (Thermo Fisher) with values averaged for dilution calculations. Suspensions were then diluted to 2×10^6^ cells/ml and 100 ul (2×10^5^ cells) were injected subcutaneously into the left flank with a 1 ml syringe and 27 gauge needle. The left flank of all mice was shaved with an electric clipper the day prior to implantation and all procedures performed under 2% isoflurane. Mice were allowed to grow for the indicated time, typically 16 (for flow cytometry) or 28 days (for evaluating metastasis) with regular monitoring and tumor measurement. Spontaneous lung metastases were identified by the presence of black surface nodules in the lung and in some cases by H&E staining. The six skin-draining LNs (left and right inguinal, axillary, and brachial nodes) were evaluated for LN metastasis by the presence of punctual black melanoma cells growing within the node and metastatic area was measured with digital caliper. LN metastatic burden is represented by summing the total LN metastatic area of all invaded nodes in each mouse. When quantifying number of LN metastases, distinct metastases in a LN were counted independently due to presumed independent seeding.

### Cancer immunotherapy models

B16F0 or LN8-1198AR cell lines were implanted into mice as described above. Seven days after implantation, mice were randomized across treatment groups. Timing and number of doses for each therapy are indicated in the respective figures. Combination ICB therapy consisted of 100 ug of aPD-L1 (BioXCell, Clone 10F.9G2) and 100 ug of aCTLA-4 (BioXCell, Clone 9H10) given intraperitoneally in 200 ul of PBS with a 31 gauge insulin syringe. Intratumoral IFNa-sensitization therapy used 10 ug of IFNa produced as described below in 25 ul of PBS delivered using a 31 gauge insulin syringe. 50 ug of IFNa-MSA was used for systemic IFNa-sensitization therapy, delivered intraperitoneally with a 31 gauge insulin syringe.

### Experimental Lung Metastasis Model

B16F0 and LN6-987AL cell lines were grown as described as above. Each cell line had previously been transduced with a lentivirus encoding for tdTomato to enable detection by a fluorescence dissection microscope. Cells were selected periodically (every ∼6 passages) in 400 ug/ml Hygromycin (Thermo Fisher) to ensure a pure population of fluorescent cells before a minimum of 2 passages without selection prior to use. Cells were prepared for injection as described above for subcutaneous implantation and diluted to 1×10^6^ cells/ml in phenol red-free DMEM. Mice were anaesthetized with 2% Isoflurane, their tails warmed in water that was warm to the touch, before injection of 200 ul of cells (2×10^5^ cells) into the lateral tail vein with a 1 ml syringe and a 27 gauge needle. Mice were then monitored and lung metastases allowed to grow for 12-18 days as indicated before euthanasia. At euthanasia, lungs bearing fluorescent cells were inflated with PBS and imaged with a fluorescence dissection microscope to enable counting of metastatic nodules. Lung bearing non-fluorescent cells were inflated with fixative and prepared for H&E staining and imaging as described below to quantify metastatic burden.

### Hematoxylin & Eosin (H&E) Staining

Lungs were inflated with 10% formalin (Sigma-Aldrich) or 4% Paraformaldehyde (Electron Microscopy Sciences), fixed overnight at 4 C, then transferred to 70% Ethanol. LNs were removed from the mouse and placed directly in 10% formalin or 4% Paraformaldehyde and fixed for approximately 4 hours before transfer to 70% ethanol. Tissues stored in 70% ethanol were then blocked, sectioned, and stained with H&E by HistoTech. Tissue sections were then imaged using an EVOS M7000 microscope (Thermo Fisher).

### Evolution of Spontaneous B16F0 and MOC2 Metastatic Models

B16F0 triad cell lines (868 lineage) were derived by implanting B16F0 cells subcutaneously as described above and allowing tumors to grow for 28-30 days. At the endpoint, a mouse with both an inguinal (tumor-draining LN) and multiple lung metastasis nodules was found. Non-tumor tissue was removed when possible from the lung and LN nodules to minimize normal cell contamination, and multiple non-necrotic regions of the primary tumor were collected and pooled. Tissues were minced with scissors in trypsin, incubated for 5 minutes at 37 C, quenched with cDMEM, and filtered through a 100 um filter, with any remaining chunks mashed with the back of a syringe. Cell suspensions were centrifuged at 250 x g, supernatant removed, and plated in an appropriate size well in cDMEM. The following day dead cells and immune cells were removed by washing the well with cDMEM. When confluent, each line was expanded over 3-5 passages before cryopreservation for downstream assays.

MOC2 lung and LN metastatic cell lines were derived through two rounds of sequential orthotopic implantation and isolation. MOC2 cells were implanted in the buccal region and grown for 2-3 weeks. At the endpoint, mice were dissected and tumor-draining cervical LNs or metastatic lung nodules were isolated. Primary tumor tissue was not collected due to recurrent fungal and bacterial contamination during culture. Tissues were minced with scissors in trypsin, incubated for 5 minutes at 37 C, quenched with cDMEM, and filtered through a 100 um filter, with any remaining chunks mashed with the back of a syringe. Cell suspensions were centrifuged at 250 x g, supernatant removed, and plated in an appropriate size well in cDMEM. The following day dead cells and immune cells were removed by washing the well with cDMEM. Second generation lines were generated by repeating the same process using the expanded first-generation LN and lung metastatic MOC2 cell lines, with resulting lines cryopreserved for downstream assays.

### Generation of Knockout Cell Lines

sgRNA sequences targeting murine Ifnar1, Ifnar2, Ifngr1, or Stat1 (Supplementary Table 1) were cloned into the PX458 plasmid for transient expression of a sgRNA, Cas9, and eGFP reporter using BbsI and Quick Ligase. Plasmids were grown in Stbl cells and isolated plasmids were confirmed by whole-plasmid sequencing (Plasmidsaurus).

For each knockout, either B16F0 and LN6-987AL cells were transiently transfected with a given PX458 plasmid using Lipofectamine 3000 (Thermo Fisher Scientific) according to the manufacturer’s protocol. Cells were transfected in 10 cm dishes with 15 ug of plasmid for 18 hours, before expansion in cDMEM for 1-3 days prior to selection. GFP+ cells were sorted using a FACSAria II with a 100 um nozzle and expanded. The transfection and sorting steps were repeated a second time for each knockout. To remove cells with Cas9 integration, a third sort was performed after overnight stimulation with universal IFNa or murine IFNy and staining for H-2Kb PE (Clone AF6-88.5), selecting for GFP-H-2Kb-cells. Cells were expanded in cDMEM and frozen for use in experiments.

### Chronic Interferon Evolution

B16F0 and YUMM1.7 cells were grown as indicated above. Cells were seeded in 6-well plates at sufficient density to reach confluency in ∼48 hours and grown in 2 ml cDMEM at 37 C and 5% CO2. Cells chronically stimulated were grown in constant 500 U/ml Universal IFNa (PBL Assay Science) and control cells without stimulated were grown at the same time. Cells were split when they reached confluency and seeded as described for 4 weeks. After 4 weeks of culture, cells were then prepared and implanted in mice as described above for the evaluation of tumor growth and metastasis.

Evolution was continued for an additional 8 weeks (total of 12) for bulk RNA-seq. After 12 weeks of evolution, cells were removed from IFNa and routinely split for one week, before being split and restimulated with 1000 U/ml Universal IFNa. 24 hours later, the cells were trypsinized, washed with cDMEM followed by PBS, then flash frozen as −80 C until RNA extraction, library preparation, and sequencing by MedGenome.

### Flow Cytometry of Cell Lines

Cells were grown as indicated above and used at the lowest passage number possible. Cells were seeded in 6-well plates 24 hours prior to analysis at proper density to reach ∼70% confluence the next day. Cells were grown in 2 ml of cDMEM at 37C and 5% CO2. Stimulation with Universal IFN-a (PBL Assay Science) was performed at 500 U/ml for 24 hours. The next day, cells were detached from plates with .5 ml Accutase for 2-5 minutes at 37C, quenched with 2 ml of cDMEM, and collected in FACS tubes. Cells were then washed with FACS Buffer and transferred to 96 well U-bottom plates for staining and analysis. Cells were stained in 100 ul of FACS Buffer with antibodies against H-2Db APC (Clone KH95), H-2Kb PE/APC/BUV395 (Clone AF6-88.5), and PD-L1 BUV737 (Clone 10F.9G2) at 1:400 dilution and anti-CD16/32 FcBlock at .5 mg/ml (BioXCell, Clone 2.4G2) for 30 minutes at 4 C in the dark. Cells were then washed twice with FACS Buffer before data was collected on a Quanteon or Penteon Instrument (Agilent).

### Tissue Processing for Flow Cytometry and ELISAs

At the indicated timepoint (typically 16 or 28 days after tumor implantation), mice were euthanized and tumor and draining lymph node (left inguinal) were collected in FACS Buffer on ice. 100-400 mg of tumor was used for subsequent processing with necrotic tissue removed when possible. Tissues were weighed, placed in a glass scintillation vial, and chopped with scissors until pieces were ∼1 mm^3^ in digestion buffer (RPMI-1640 with 1 mg/ml Collagenase 4 and .1 mg/ml DNase I). Tissues were incubated on a magnetic stir plate at 37 C for 10 minutes for lymph nodes and 20 minutes for tumors. Digestions were quenched with an equal volume of ice-cold RPMI-1640 with 10% FBS and tissues were maintained at 4 C for the remainder. Digested tissues were then filtered through a 70 um strainer, remaining debris mashed with the back of a syringe plunger, and washed with FACS buffer. For cytokine stimulation and ELISA, cell suspensions were pelleted at 350 x g for 5 minutes, supernatant aspirated, and resuspended in FACS buffer before aliquoting into plates for subsequent assays. For tumor microenvironment profiling, cell suspensions were pelleted at 350 x g for 5 minutes, supernatant aspirated, and resuspended in 30% Percoll solution. The cell-Percoll solution was then layered on top of 80% Percoll in a 15 ml conical tube and centrifuged for 15 minutes at 400 x g with 0 brake. The immune-enriched live cell layer at the interface of the two layers was aspirated and washed with FACS buffer before aliquoting into plates and proceeding with flow cytometry staining.

### Flow Cytometry of Tissues

Single cell suspensions of lymph nodes (half of a lymph node) or tumors (100 mg of tissue pre-Percoll isolation for phenotyping, 25 mg of tissue for stimulation) were used for staining and analysis. For interferon detection, cells were cultured in cRPMI for 12 hours with Cell Activation Cocktail with Brefeldin A (40.5 uM PMA and 669.3 uM ionomycin, 2.5 mg/ml Brefeldin A), supplemented with Monensin (2 uM).

Cells were stained with Live/Dead Blue at 1:500 in 100 ul of FACS Buffer, then washed twice with FACS Buffer. Surface epitopes sensitive to fixation were then stained in 100 ul of FACS Buffer with anti-CD16/32 FcBlock at .5 mg/ml and Brilliant Stain Buffer (final 10%) in the dark at 4C for 30 minutes, then washed twice with FACS Buffer. Cells were then fixed in 100 ul with either the eBiocience Foxp3/Transcription Factor Fixation buffer (immune profiling) or the BD Cytofix/Cytoperm Fix/Perm buffer (interferon detection) for 30 minutes at 4 C in the dark. After fixation, the cells were washed twice with eBiocience Foxp3/Transcription Factor Wash buffer (immune profiling) or BD Cytofix/Cytoperm Perm/Wash buffer (interferon detection). Non-fixation sensitive surface epitopes and intracellular epitopes were then stained in 100 ul of the respective wash buffer with anti-CD16/32 FcBlock at .5 mg/ml and Brilliant Stain Buffer (final 10%) at 4 C in the dark overnight (14-18 hours)^85^. Cells were washed with the respective wash buffer twice, resuspended in FACS Buffer supplemented with AccuCount Fluorescent Particles for normalization. Data was acquired using a Cytek Aurora 5-laser instrument (Cytek Biosciences) with unmixing performed in SpectroFlo (Cytek Biosciences, Version 3.3.0) before analysis in FlowJo (Becton, Dickinson & Company, Version 10.10.0). All samples were run within 72 hours of staining to minimize sample breakdown. Antibody panels including marker, fluorophore, clone, dilution, staining step, and unmixing sample type are included in Supplementary Table 2 (Interferon Panel), Table 3 (Lymphoid Panel), and Table 4 (Myeloid Panel).

### NK Cell Cytotoxicity Assay

Splenocytes were isolated from C57BL/6 mice by disrupting the spleens against a 70 um filter on a 50 ml tube with repeated washing with FACS buffer. Cell suspensions were then pelleted at 350 xg for 5 minutes, resuspended in ACK lysis buffer for 2 minutes to remove red blood cells, and quenched with PBS. Cell suspensions were again pelleted at 350 xg for 5 minutes before proceeding to NK cell isolation with the EasySep Mouse NK Cell Isolation Kit (Stem Cell Technologies) according to the manufacturer protocol. Tumor cells were detatched with Accutase and resuspended to 2×10^6^ cells/ml in HBSS with 2% FBS and 5 uM Calcein-AM (Thermo Fisher). Cells were stained for 1 hour at 37 C, then washed twice with HBSS with 2% FBS. NK and tumor cells were then mixed at a 10:1 ratio (10^5^ NK cells:10^4^ tumor cells) in a 96 well U-bottom plate in 200 ul of RPMI-1640 supplemented with 10% FBS, 2 mM L-Glutamine, 15 mM HEPES, 1% MEM NEAA and 50 uM Beta-Mercaptoethanol. 10^4 tumor cells alone or with 10% Tween-20 were also cultured in 200 ul of complete RPMI-1640 as spontaneous lysis and maximum lysis controls.

After 4 hours of culture in an incubator at 37 C and 5% CO2, plates were centrifuged at 350 xg for 5 minutes, and supernatants transferred to a new plate. Plates were read on a Spark fluorescence plate-reader (Tecan) with an excitation of 495 nm and emission of 515 nm. Specific lysis was calculated as % Specific Lysis = 100 x (Sample – Spontaneous Lysis) / (Tween-20 – Spontaneous Lysis).

### Tumor-T cell Co-Culture

Splenocytes were isolated from C57BL/6 or OT-I mice by disrupting the spleens against a 70 um filter on a 50 ml tube with repeated washing with FACS buffer. Cell suspensions were then pelleted at 350 xg for 5 minutes, resuspended in ACK lysis buffer for 2 minutes to remove red blood cells, and quenched with PBS. Cell suspensions were again pelleted at 350 xg for 5 minutes before proceeding to CD8 T cell isolation with the EasySep Mouse CD8+ T Cell Isolation Kit (Stem Cell Technologies) according to the manufacturer protocol. B16F0 or B16F0 expressing cytoplasmic ovalbumin (cOVA) were detatched with Accutase and washed with cDMEM. CD8 T cells and tumor cells were then mixed at a 1:1 ratio (10^5^ CD8 T cells:10^5^ tumor cells) in a 96 well U-bottom plate in 200 ul of RPMI-1640 supplemented with 10% FBS, 2 mM L-Glutamine, 15 mM HEPES, 1% MEM NEAA and 50 uM Beta-Mercaptoethanol. Positive control wells were supplemented with 1000 U/ml universal IFNa. The cells were cultured overnight (24 hours) at 37 C and 5% CO2. Cells were pelleted at 350 x g and stained with Live/Dead Blue at 1:500 in FACS buffer for 15 minutes, then washed twice with FACS buffer. Cells were stained in 100 ul of FACS Buffer with antibodies against H-2Db APC (Clone KH95), H-2Kb PE (Clone AF6-88.5), PD-L1 BUV737 (Clone 10F.9G2), CD8a Spark Blue 574 (Clone 53-6.7), and anti-CD16/32 FcBlock at .5 mg/ml (BioXCell, Clone 2.4G2) for 30 minutes at 4 C in the dark. Cells were then washed twice with FACS Buffer before data was collected on a Quanteon or Penteon Instrument (Agilent).

### Cytokine Measurement by ELISA

Mice were implanted with B16F0 or LN8-1198AR cell lines as described above, or DMEM in the case of naïve mice, and allowed to grow for 28 days. At the end point, the draining lymph node (left inguinal lymph node) and a segment of the tumor or flank skin were dissected, weighed, processed into single cell suspensions as described above. Half of a lymph node or 25 mg of tumor/skin tissue was resuspended in cRPMI and cultured for 4 hours with Cell Activation Cocktail without Brefeldin A (40.5 uM PMA and 669.3 uM ionomycin). Samples were centrifuged to pellet cells and culture supernatant flash frozen in 50 ul aliquots. Plates were stored at −80 C for up to two weeks before thawing at room temperature and completion of ELISAs for mouse IFNg (Biolegend), IFNa (PBL Assay Science), and IFNb (PBL Assay Science) according the manufacturer instructions.

### Interferon Production, Purification, and Activity Assay

The gene encoding recombinant IFN-aA/D(BglII)^56^ was cloned into a pD649 mammalian expression vector (ATUM DNA 2.0) containing an HA secretion signal peptide and C-terminal 6-His tag. Extended half-life versions of IFNa were generated by fusing the gene encoding mouse serum albumin to either the N or C-terminus of the IFN-aA/D(BgIII) construct. Sequences of each expression construct are detailed in Supplementary Table 5. Proteins were expressed in Expi293 cells for 3 days according to manufacturer instructions. Proteins were purified via Ni^2+^ affinity chromatography followed by fast protein liquid chromatography via a Superdex 75 increase column (Cytiva) for untagged protein or Superex 200 increase column (Cytiva) for MSA-tagged proteins, equilibrated with PBS (Gibco).

For pSTAT signaling assays, B16F0 cells were grown and detached from flasks with TrypLE Express for 3 min at 37 C. Cells were plated at 1×10^5^ cells per well in a flat-bottom 96 well plate. The next day, cells were stimulated with interferon at the indicated concentration in serum-free DMEM for 20 min at 37 C before detachment with TrypLE Express. Without washing, cells were fixed with 16% paraformaldehyde for 10 min at 37 C while shaking. Cells were washed twice with FACS buffer before permeabilization with ice-cold 100% methanol at −80 C overnight. The cells were thawed and washed twice with FACS buffer before staining with anti-human/mouse pSTAT1 conjugated to AF647 in FACS buffer for 1 hour at room temperature while shaking at 500 RPM. Cells were washed twice then analyzed by flow cytometry on a Beckman-Coulter Cytoflex-S.

### RNA Sequencing of Cell Lines

Samples of newly derived cell lines were collected for sequencing at the time of first cryopreservation, typically 2-5 passages after isolation. Cells were detached with .25% Trypsin-EDTA, quenched with cDMEM, and pelleted by centrifuge at 250 xg for 5 minutes. Prior to cryopreservation, an aliquot of 2-10×10^6^ cells were pelleted in a DNA low bind microcentrifuge tube and washed with PBS. Supernatant was aspirated and cell pellets flash frozen at −80 C until library preparation. Samples were transferred to MedGenome where RNA-sequencing was performed. Bulk RNA was extracted from the cell pellets, quality confirmed by TapeStation (Agilent), before library preparation with the NEB mRNA polyA kit.

Samples were pooled and sequenced on a NovaSeq 6000 (Illumina) to a minimum read depth of 20M PE100 reads.

### Analysis of Cell Line RNA-seq

Counts data from previously sequenced B16F0 or LN metastatic cell lines was normalized using edgeR and differentially expressed genes identified by comparing late-stage cell lines to parental B16F0. Differentially expressed genes that were upregulated in late-stage cell lines were plotted. Gene names were extracted and annotated as ISGs using Interferome^39^, set to default settings and searching genes involved in type I interferon processes in mouse samples.

B16F0 cells evolved in chronic IFNa were analyzed using DESeq2 using a design that accounts for both whether cells were evolved in IFNa and whether after wash-out they were re-stimulated, as well as the interaction between these two terms. Comparisons were then extracted for stimulated on non-evolved cells and cells evolved against those not, without re-stimulation. Differentially expressed genes in evolved cells compared to non-evolved cells were used to generate a heatmap, with Z-scores generated independently for basal and re-stimulated cells to account for scale differences.

B16F0 triad cell lines (originating from primary tumor, LN metastasis, or lung metastases) were analyzed by converting the counts matrix to log2(TPM+1) values for plotting. A combined list of IFNa and IFNb responsive genes were used for plotting. Genes were annotated in up in LN or lung by taking the difference between log2(TPM+1) values for LN and lung samples.

### Analysis of Human RNA-seq

Bulk RNA-seq of leukocytes (CD45+ CD31-FAP-EPCAM-) and tumor cells (EPCAM-CD45-CD31-FAP-) were accessed from GEO under accession code GSE113839^25,79^. Log2(TPM+1) values for all type I interferon genes were summed to calculate type I interferon scores and remaining single gene values were taken for remaining interferons. Patient characteristics, including nodal involvement, for the patients included in the analysis are available in Supplementary Table 7.

Paired primary tumor and metastasis bulk RNA-seq of breast cancer patients was obtained from GEO under accession code GSE209998. Log2(FoldChange) gene expression values were calculated for each patient comparing their metastasis to their primary tumor, averaging across all genes in each gene set in Supplementary Table 6 to compute metagene scores. Solid organ metastases include patients with liver, ovarian, or lung metastases.

BMS038 bulk RNA-seq data of pre- and on-therapy melanoma patients receiving ICB was accessed from GEO under accession code GSE91061. Metagene scores were calculated for the gene sets in Supplementary Table 6 using pre- or on-therapy samples for each patient. Survival curves were stratified by the original responder/non-responder annotation and their pre-therapy interferon score using the survminer package (Version 0.5.1).

### Quantification and Statistical Analyses

Statistical analyses were performed either in Prism (Version 10.5.0) or R (Version 4.5.1). Individual data points are shown when possible, with bar heights or horizontal lines corresponding to means. In all figures, statistical significance annotations correspond as follows: * p < .05, ** p < .01, *** p < .001, **** p < .0001. Differences between categorial variables was assessed either by Fischer’s Exact Test or Chi-squared Test depending on the sample size. Comparisons between two groups were computed using either a two-tailed Welch’s t test or in the case of non-normal data (such as lung metastasis counts) a two-tailed Mann Whitney test. In the case of comparing more than three groups, a one-way ANOVA was used, utilizing the Brown-Forsythe and Welch ANOVA test with Dunnet’s T3 multiple comparisons test or a Kruskal-Wallis test with Dunn’s multiple comparison test depending on the normality of the data. Longitudinal data or grouped data (such as tumor growth curves and flow cytometry of knockout tumors) was analyzed using a two-way ANOVA and Tukey’s multiple comparisons test. Additional details for specific comparisons are reported in the respective figure legends.

## Data and Materials Availability

LN-metastatic cell lines previously derived from B16F0 are available from Cancer Tools^84^. Knockout cell lines or newly derived metastatic cell lines are available from the lead contact upon request and completion of a materials transfer agreement with Stanford University.

RNA-seq of B16F0 LN metastatic cell lines has previously been made available on GEO under accession number GSE117529. Additional RNA-seq of knockout cell lines and cells evolved under chronic IFNa will be deposited in GEO at the time of publication. RNA-seq of primary and paired metastases from breast cancer patients was obtained from GEO under accession number GSE209998. RNA-seq of pre/on-therapy samples from BMS038 was obtained from GEO under accession number GSE91061. RNA-seq of leukocytes and tumor cells isolated from HNSCC primary tumors was accessed from GEO under accession number GSE113839. Raw and processed 4T1 metastasis barcode data was obtained from Simon Knott^13^.

## Author Contributions

CBB and NERF conceptualized the study and designed all experiments. CBB performed all experiments, flow cytometry, and analyses except derivation of new MOC2 cell lines and protein production. ML assisted with flow cytometry and MOC2 experiments. GER produced/purified proteins and assisted with therapeutic studies. NERF and KCG supervised the study. CBB and NERF wrote the manuscript with feedback from all authors.

## Acknowledgements

We thank Amanda Kirane and Lisa Nichols for helpful discussion. Flow cytometry and cell sorting were performed using instruments in the Stanford Shared FACS Facility including an instrument obtained using NIH S10 Shared Instrument Grant S10RR025518-01. This work was supported by NIH grant DP2 AI177915 (NERF), a Stanford Cancer Institute Innovation Award (NERF), NIH grant R01 AI51321 (KCG), the Mark Foundation (KCG), the Howard Hughes Medical Institute (KCG), and the Parker Institute for Cancer Immunotherapy (KCG). CBB is supported by an Arc Institute Fellowship and NCI F31 CA301857. GER was funded by the Parker Institute for Cancer Immunotherapy.

---

**Supplementary Figure 1.**
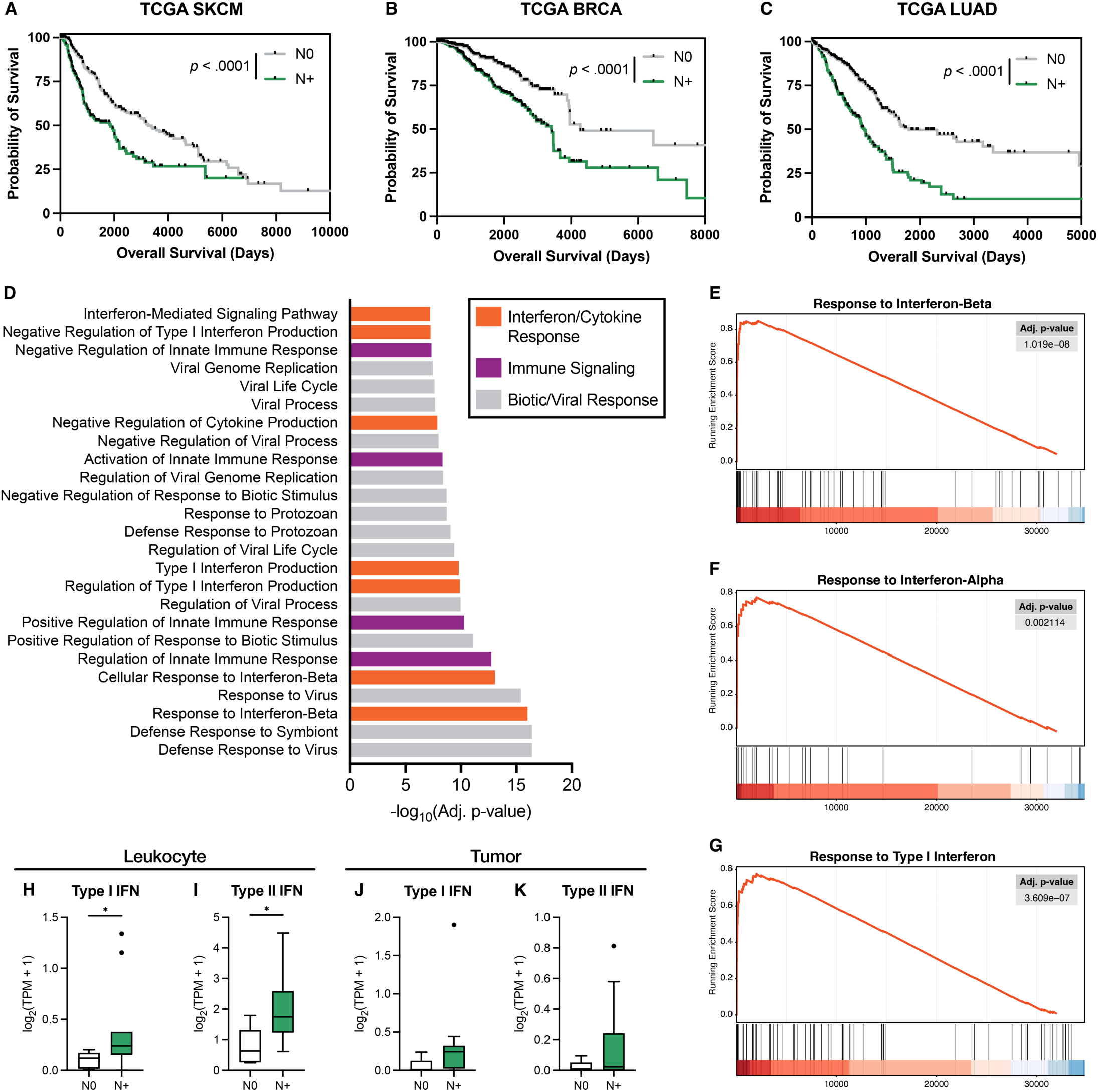
(A) SKCM, (B) BRCA, and (C) LUAD TCGA dataset survival curve plots stratified according to N0 or N+ status. (D) Ranked GO terms from analysis comparing late-stage LN metastatic cell lines to parental cell line. (E-G) GSEA plots for type I interferon response terms for late-stage LN metastatic cell lines. Type I and II interferon gene expression counts from (H, I) bulk sorted leukocyte and (J, K) tumor cells from HNSCC patients with or without nodal involvement.

**Supplementary Figure 2.**
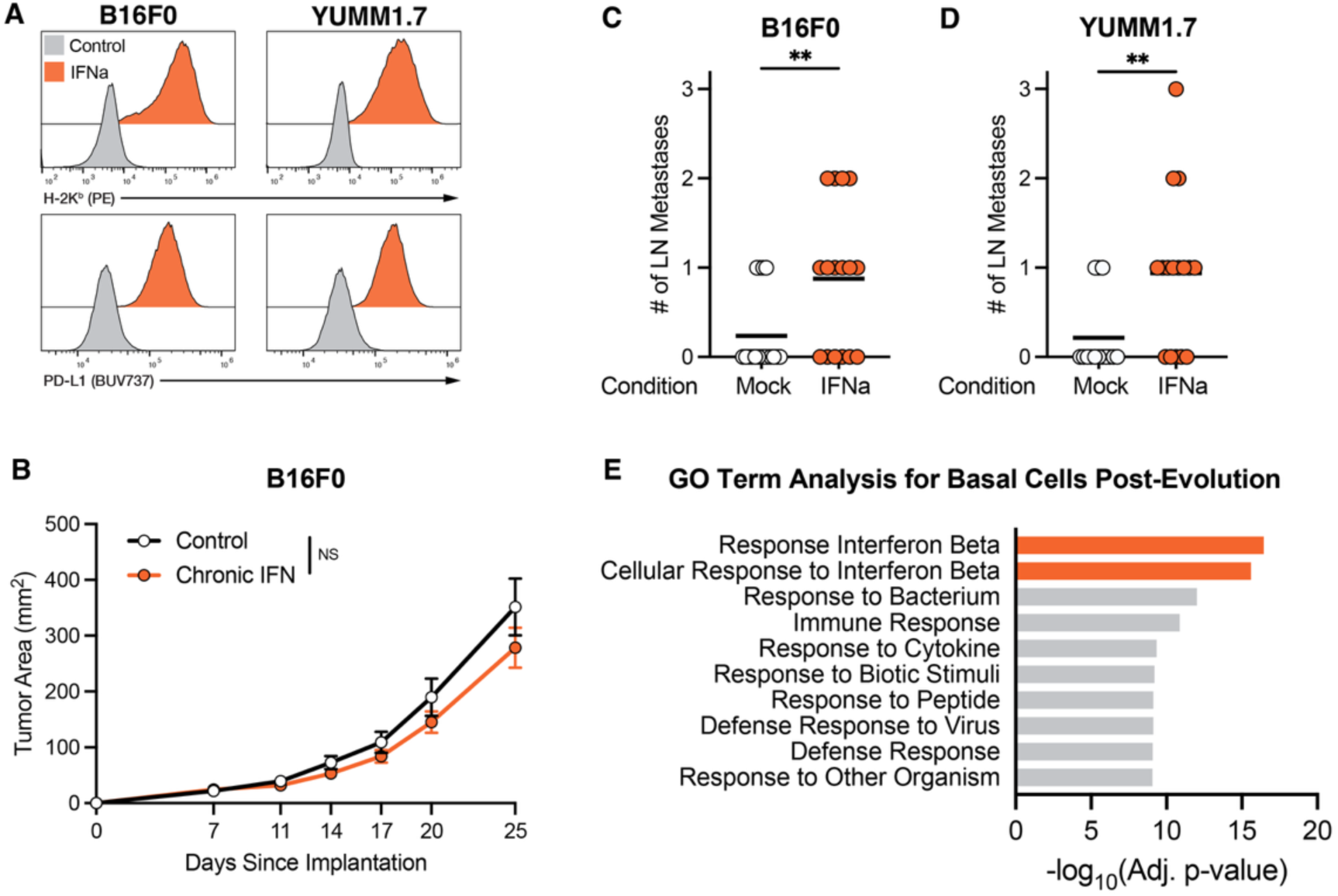
(A) Representative histograms of H-2Kb and PD-L1 expression for mock or chronic IFNa evolved cell lines at the end of four weeks of culture. (B) Tumor growth curves for B16F0 mock evolved or chronic IFNa evolved cells implanted in mice. (C) B16F0 and (D) YUMM1.7 LN metastasis counts for mock or chronic IFNa evolved cell lines in mice. (E) Ranked GO terms from analysis comparing chronic IFNa evolved cells to mock evolved cells after wash-out without re-stimulation.

**Supplementary Figure 3.**
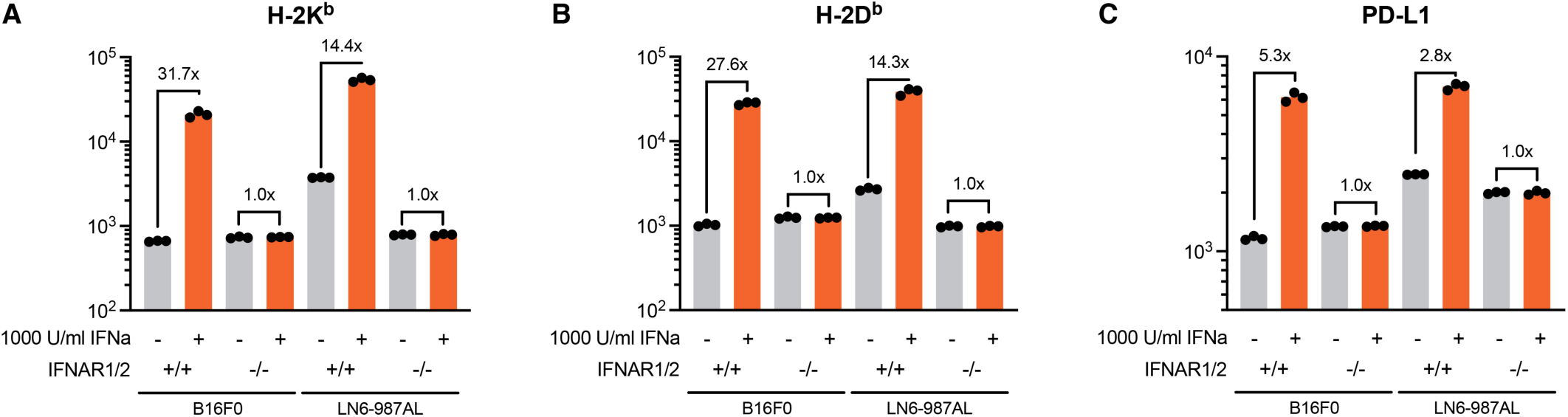
(A) H-2Kb, (B) H-2Db, and (C) PD-L1 geometric MFI on B16F0 and LN6-987AL wildtype or IFNAR1/2 double-knockout cells upon stimulation with IFNa, confirming loss of signaling.

**Supplementary Figure 4.**
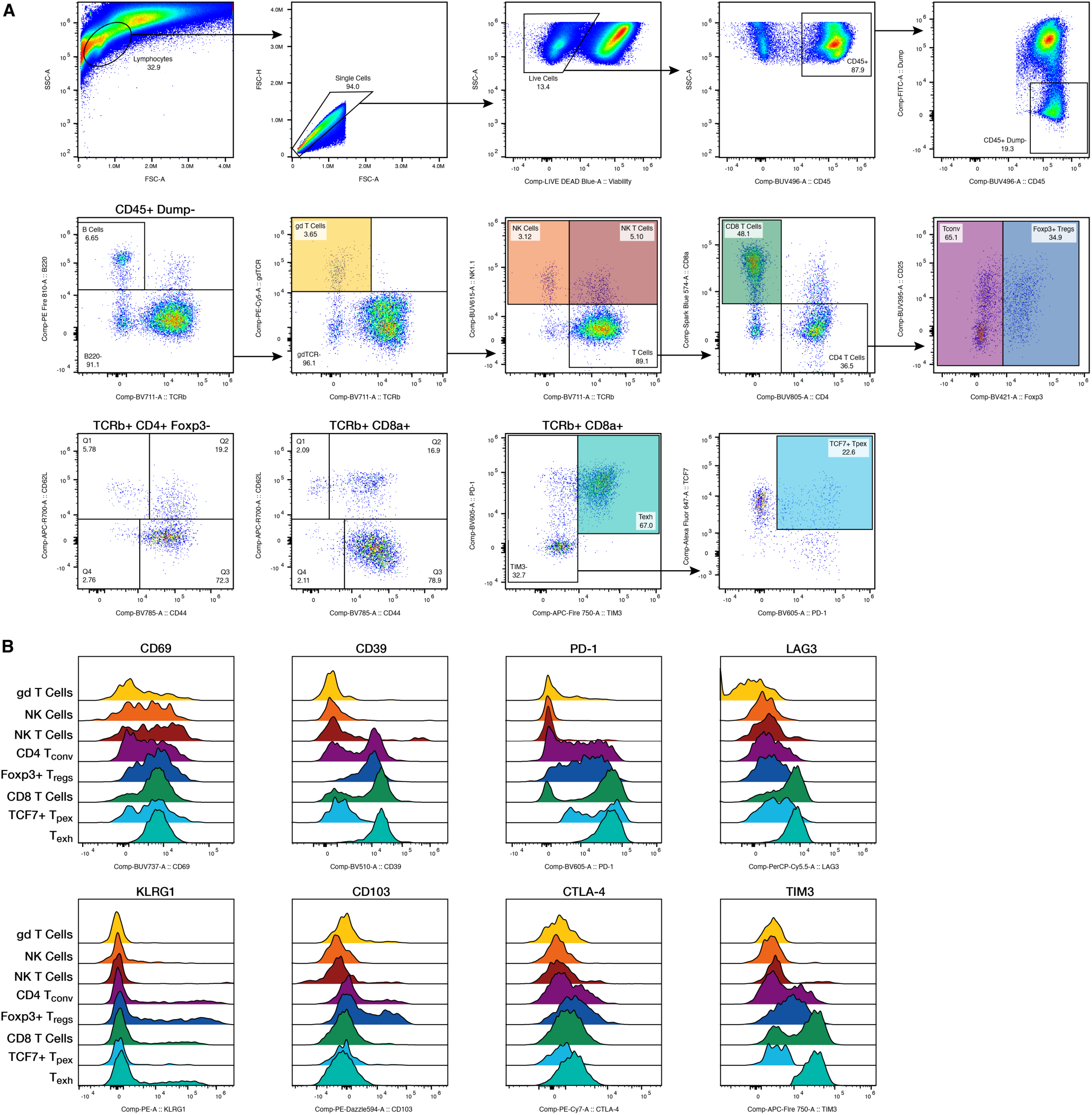
Gating scheme for identification of lymphoid cell populations. (B) Histograms depicting functional marker expression for each gated cell population in (A), normalized to mode.

**Supplementary Figure 5.**
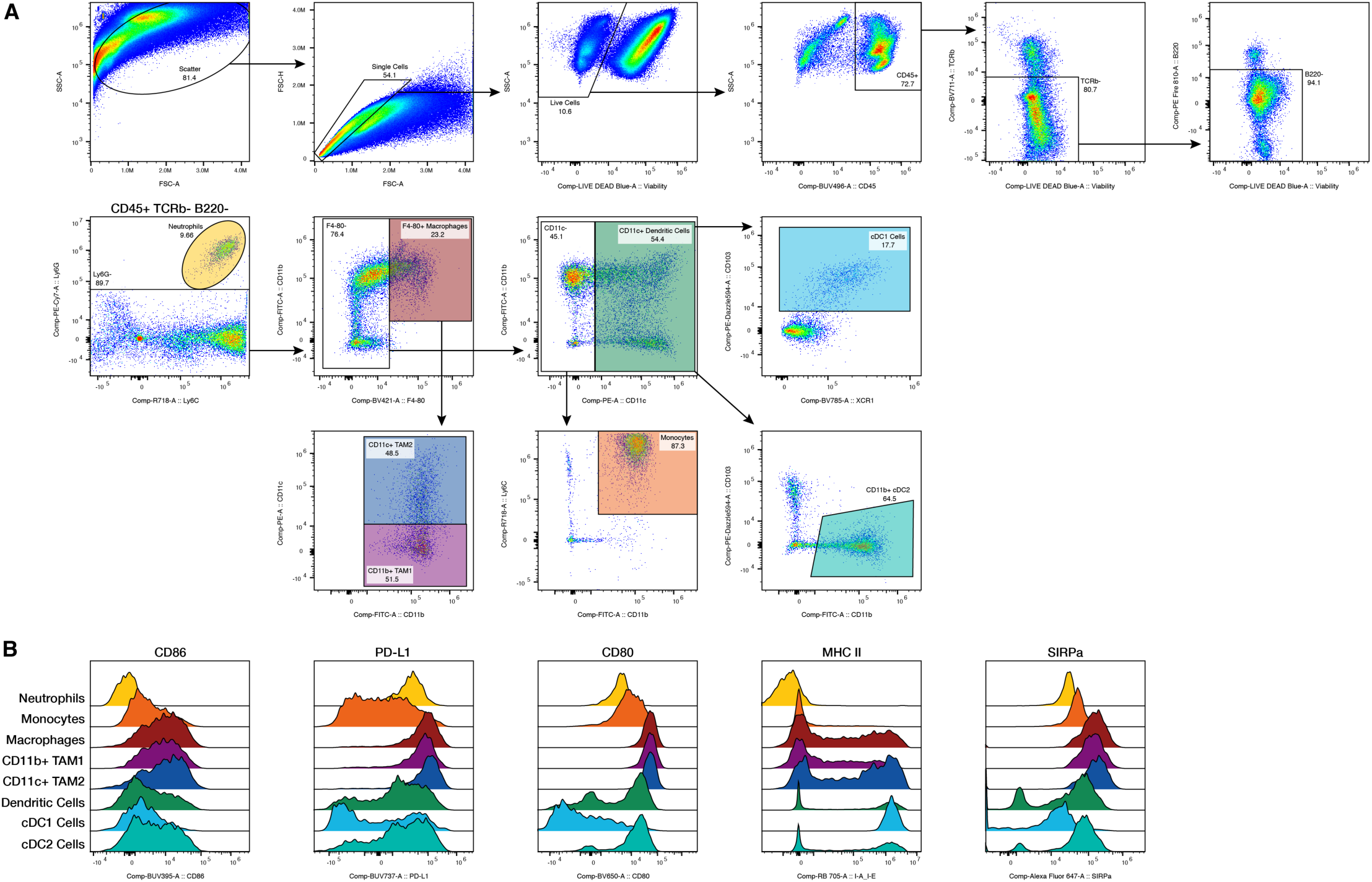
Gating scheme for identification of myeloid cell populations. (B) Histograms depicting functional marker expression for each gated cell population in (A), normalized to mode.

**Supplementary Figure 6.**
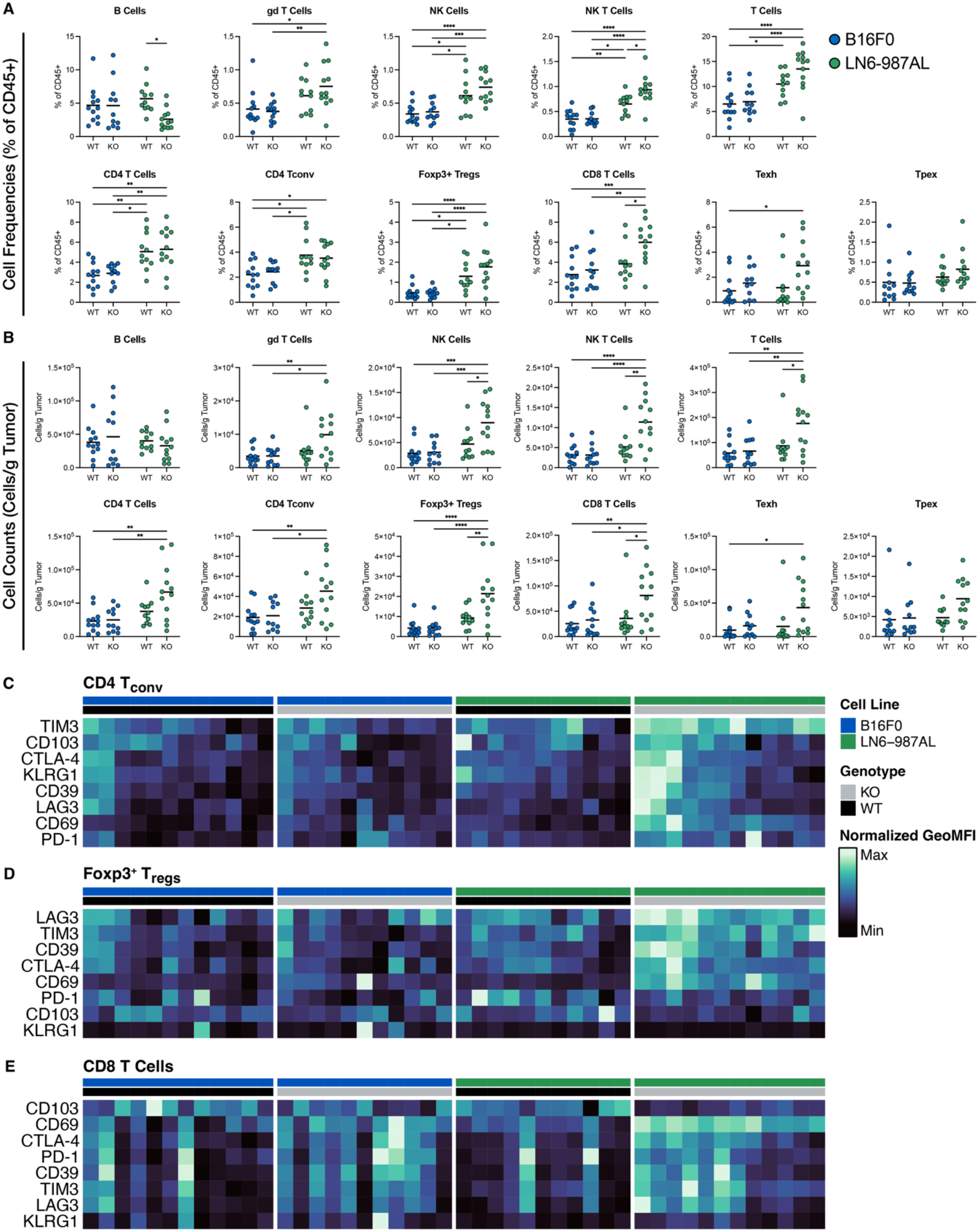
(A) Cell frequencies (% of CD45+ cells in the tumor) and (B) cell counts normalized to tumor mass for lymphocyte cell populations in B16F0 or LN6-987AL tumors according to wildtype or IFNAR1/2 knockout status. (C) CD4 Tconv, (D) Foxp3+ Treg, and (E) CD8 T cell functional immune cell marker geometric MFIs, scaled within each cell population.

**Supplementary Figure 7.**
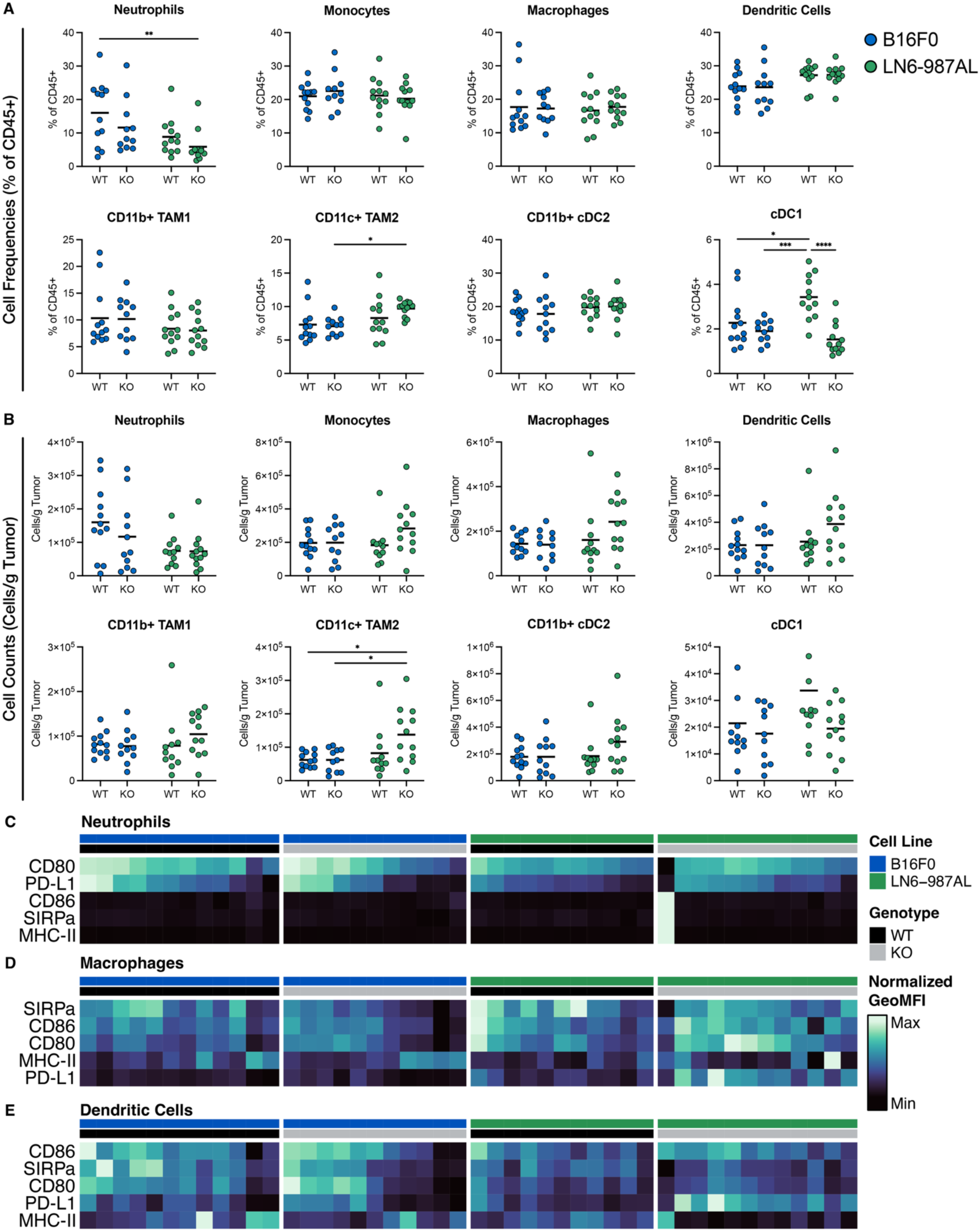
(A) Cell frequencies (% of CD45+ cells in the tumor) and (B) cell counts normalized to tumor mass for myeloid cell populations in B16F0 or LN6-987AL tumors according to wildtype or IFNAR1/2 knockout status. (C) Neutrophil, (D) Macrophage, and (E) Dendritic Cell functional immune cell marker geometric MFIs, scaled within each cell population.

**Supplementary Figure 8.**
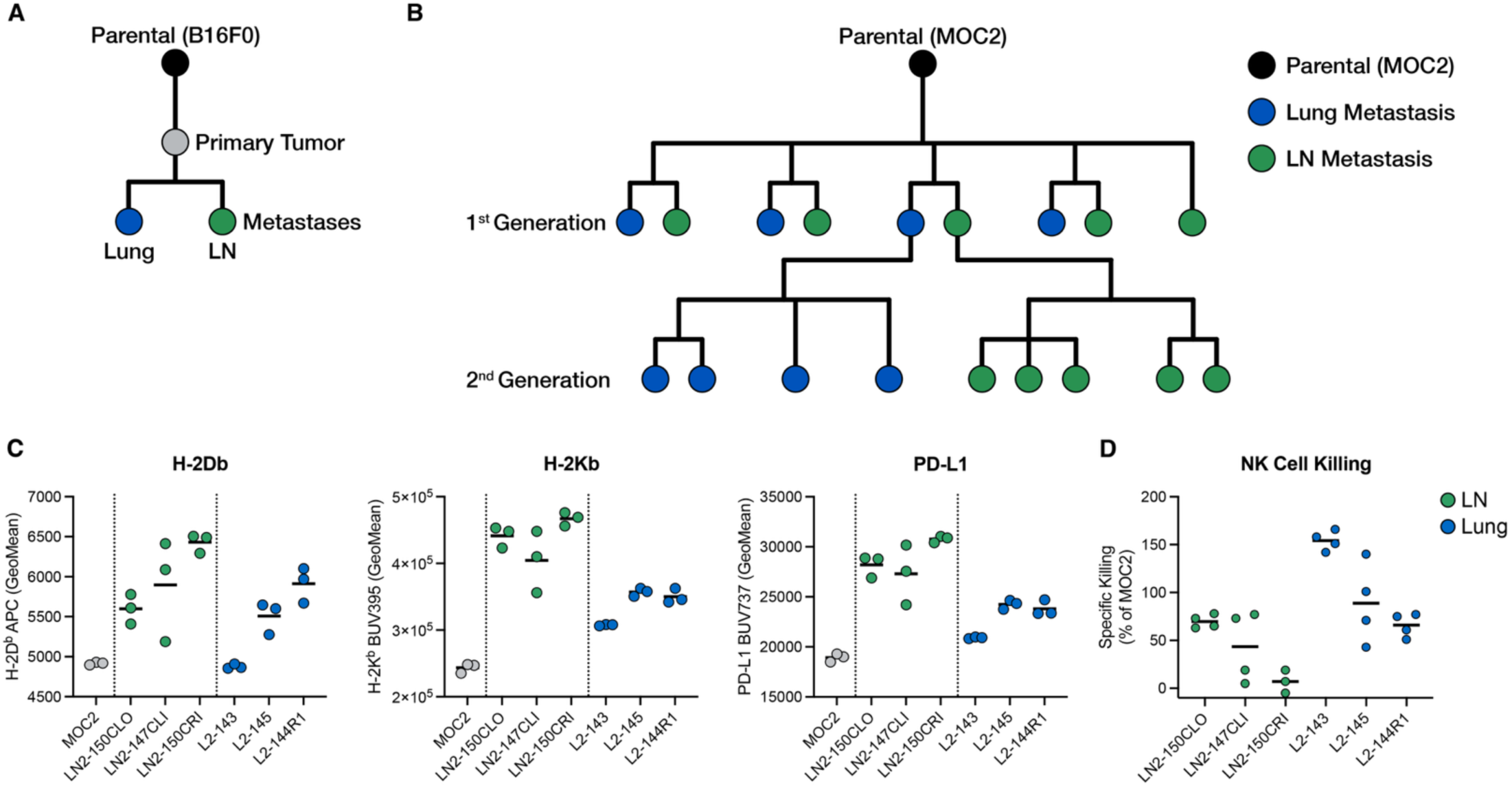
(A) MOC2-derived cell line phylogeny for LN and lung metastatic branches. Short branches indicate cell lines derived from the same mouse while long branches indicate individual mice. (B) B16F0-derived 868 primary tumor, lung metastasis, and LN metastasis phylogeny. (C) H-2Db, H-2Kb, and PD-L1 geometric MFIs for each MOC2-derived cell line evaluated at baseline in culture. (D) Corresponding in vitro NK cell specific killing of each cell line expressed as a percent of control MOC2 killing.

**Supplementary Figure 9.**
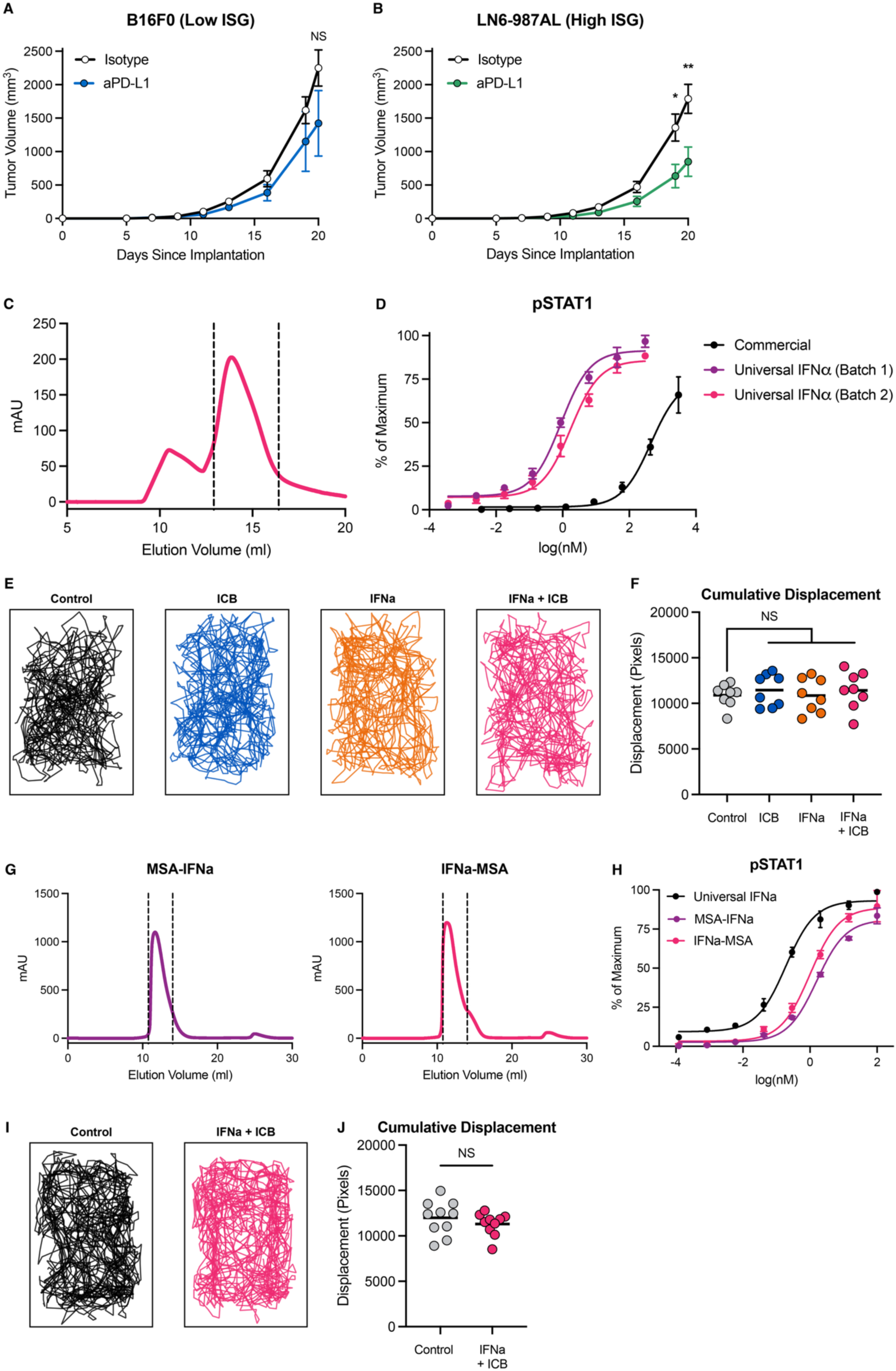
(A) B16F0 and (B) LN6-987AL tumor growth curves receiving anti-PD-L1 monotherapy. (C) FPLC trace of universal IFNa production, with region between vertical lines corresponding to fraction collected for further testing. (D) pSTAT1 B16F0 dose-response curves for universal IFNa and commercial protein by flow cytometry. (E) Individual mouse traces and (F) cumulative displacement for mice treated with intratumoral monotherapy or combination therapy, evaluated 24 hours after second dose of each therapy. (G) FPLC traces for N and C-term MSA-fused extended half-life universal IFNa. Region between each pair of vertical lines was collected for further testing. (H) pSTAT1 B16F0 dose-response curves for N/C-term constructs compared to previously produced universal IFNa. (I) Individual mouse traces and (J) cumulative displacement for mice treated with systemic combination therapy, evaluated 24 hours after second dose of each therapy.

